# Species Tree Inference on Data with Paralogs is Accurate Using Methods Intended to Deal with Incomplete Lineage Sorting

**DOI:** 10.1101/498378

**Authors:** Zhi Yan, Megan L. Smith, Peng Du, Matthew W. Hahn, Luay Nakhleh

**Affiliations:** Department of Computer Science, Rice University, 6100 Main Street, Houston, TX 77005, USA; Department of Biology and Department of Computer Science, Indiana University, 1001 East Third Street, Bloomington, IN 47405, USA; Department of BioSciences, Rice University, 6100 Main Street, Houston, TX 77005, USA

**Keywords:** Multispecies coalescent, incomplete lineage sorting, gene duplication and loss, orthology, paralogy

## Abstract

Many recent phylogenetic methods have focused on accurately inferring species trees when there is gene tree discordance due to incomplete lineage sorting (ILS). For almost all of these methods, and for phylogenetic methods in general, the data for each locus is assumed to consist of orthologous, single-copy sequences. Loci that are present in more than a single copy in any of the studied genomes are excluded from the data. These steps greatly reduce the number of loci available for analysis. The question we seek to answer in this study is: What happens if one runs such species tree inference methods on data where paralogy is present, in addition to or without ILS being present? Through simulation studies and analyses of two large biological data sets, we show that running such methods on data with paralogs can still provide accurate results. We use multiple different methods, some of which are based directly on the multispecies coalescent (MSC) model, and some of which have been proven to be statistically consistent under it. We also treat the paralogous loci in multiple ways: from explicitly denoting them as paralogs, to randomly selecting one copy per species. In all cases the inferred species trees are as accurate as equivalent analyses using single-copy orthologs. Our results have significant implications for the use of ILS-aware phylogenomic analyses, demonstrating that they do not have to be restricted to single-copy loci. This will greatly increase the amount of data that can be used for phylogenetic inference.

---

Species tree inference often requires us to account for the fact that the evolutionary histories of different loci can disagree with each other, as well as with the phylogeny of the species. The reasons for this incongruence include biological causes such as incomplete lineage sorting (ILS) and introgression (broadly interpreted to include all biological processes involving genetic exchange), as well as technical causes such as the misidentification of paralogs as orthologs (“hidden paralogy”; Doolittle and Brown, 1994).

The inference of phylogenies can be carried out by concatenating all loci together or by treating each locus separately (reviewed in Bryant and Hahn, 2020). While concatenation ignores incongruence, gene tree-based methods allow each locus to take on its own topology. Some gene tree-based methods rely on a model for how these trees evolve within the species phylogeny (in addition to probabilistic models of sequence evolution on the gene trees). The multispecies coalescent (MSC) (Hudson, 1983; Takahata, 1989; Rannala and Yang, 2003; Degnan and Rosenberg, 2009) has emerged as the most commonly employed model of such gene genealogies. Indeed, in the last two decades a wide array of methods and computer programs have been developed for species tree inference under the MSC; see (Liu et al., 2009; Knowles and Kubatko, 2011; Nakhleh, 2013; Liu et al., 2015) for recent reviews and surveys of these methods. Other gene tree-based methods are inspired by the MSC, but do not rely explicitly on this model (e.g., Mirarab et al., 2014). In either case, the goal is for the methods to be robust to incongruence caused by ILS.

Regardless of the method being employed, the inference of species trees usually assumes that the data consist of only orthologous sequences. Indeed, most phylogenetic methods require the identification of orthologs; see Smith and Hahn (2021b) for a review of methods that do not require orthologs. As a result of the common requirement of orthologous loci, before such inference methods are applied to a phylogenomic dataset paralogs must be identified and removed from the data. One common approach for removing paralogs is to use graph-based methods to identify homologous gene families, and then to use those gene families present in exactly a single copy in each sampled genome for phylogenetic inference (e.g., Li et al., 2003). Another approach is to use branch-cutting methods to extract orthologs from larger gene families (e.g., Yang and Smith, 2014). Neither of these two approaches guarantees that the resulting data set includes only orthologous sequences (Koonin, 2005). Furthermore, restricting the data to single-copy genes—which is by far the most common practice in the community—means that much of the data must be excluded from the analysis. In particular, as more species are sampled, the frequency of genes that are present in single-copy across all species will decrease (Emms and Kelly, 2018).

Paralogous sequences are often modeled by a process of gene duplicaton and loss (GDL) (Boussau et al., 2013). This process can also produce incongruence, as every duplication event adds a single branch not found in the species tree (losses cannot generate incongruence). Although the MSC generates a distribution of gene trees due to ILS, it is likely that GDL models induce a distribution that differs from this. An obvious way to handle data sets where ILS and GDL could have simultaneously acted on gene families is to employ models of gene evolution that go beyond the MSC in order to incorporate GDL as well. Indeed, such models are beginning to emerge (Rasmussen and Kellis, 2012; Li et al., 2020). However, the more complex the models of gene family evolution, the more computationally prohibitive statistical inference under these models becomes (Du and Nakhleh, 2018), rendering their applicability infeasible except for very small data sets in terms of the number of species and gene families.

Given that much progress in terms of accuracy and computational efficiency has been made on gene tree-based, ILS-aware species tree inference methods, we ask in this paper the following question: are these inference methods robust to the presence of paralogs in the data? If they are, then the reach of gene tree-based inference methods is significantly extended and the exclusion of paralogous loci from phylogenomic data sets is deemed unnecessary, thus providing more signal for the inference task. To answer this question, we study the performance of five species tree inference methods, all of which use gene trees as the input data: The maximum pseudo-likelihood method of Yu and Nakhleh (2015) as implemented by the function `InferNetwork_MPL` in PhyloNet (Wen et al., 2018), ASTRAL-III (Zhang et al., 2018), NJ_st_ (Liu and Yu, 2011), ASTRAL-Pro (Zhang et al., 2020), and FastMulRFS (Molloy and Warnow, 2020). The latter two methods were developed with paralogs in mind, and so should serve as a good baseline for comparison to the MSC-inspired methods.

To test these methods, we use both simulated and real data. We simulate across a wide range of GDL rates and levels of ILS, and use two genome-scale empirical datasets with thousands of loci that contain branches with very different levels of discordance. We also sample the gene family data in multiple ways, in all cases finding that the inferences made by all methods are quite accurate, and are mostly identical to the accuracy of the inferences when using only single-copy orthologs. Particularly striking is the finding that these methods infer very accurate species trees when all gene tree incongruence is due to GDL, and ILS is not a factor. We find that gene tree estimation error affects the methods’ performances at a similar, or even higher, level than ILS. We also find that methods designed specifically to take GDL into account, namely ASTRAL-Pro and FastMulRFS, do not generally have higher accuracy than the other methods. Overall, our results support the use of approaches that account for gene tree incongruence, regardless of its causes.

## Methods

### Species tree inference methods

For species tree inference, we use five different methods. The first three assume that the input data come from single-copy genes:

- The maximum pseudo-likelihood inference function `InferNetwork_MPL` in PhyloNet, which implements the method of Yu and Nakhleh (2015). This method amounts to running MP-EST (Liu et al., 2010) when restricted to trees with no reticulations.
- ASTRAL-III (Zhang et al., 2018), Version 5.6.3.
- NJ_st_ (Liu and Yu, 2011).

While the maximum likelihood method of Yu et al. (2014) as implemented by the InferNetwork_ML function in PhyloNet (Wen et al., 2018) is relevant here, it is much more computationally demanding than maximum pseudo-likelihood, so we chose not to run it.

For comparison, we also use two methods that were designed specifically with paralogs in mind:

- ASTRAL-Pro (Zhang et al., 2020).
- FastMulRFS (Molloy and Warnow, 2020).

For the sake of conclusions that we draw from this study, it may be helpful to highlight the differences between these methods. `InferNetwork_MPL` optimizes a pseudo-likelihood function that is derived based on the assumptions of the MSC. This function is very different, for example, from a likelihood function based on a model of gene duplication and loss (Arvestad et al., 2009). Therefore, its accuracy in inferring species trees from data with paralogs reflects directly on the performance of MSC-based methods on such data. None of the other four methods make direct use of the MSC, though ASTRAL, ASTRAL-Pro, and NJ_st_ have all been shown to be statistically consistent under the MSC, at least when both gene lengths and the number of genes go to infinity. Their accuracy on data with paralogs therefore reflects the suitability of these methods, rather than the MSC itself, for analyzing such data. Finally, ASTRAL and ASTRAL-Pro are statistically consistent under the MSC and GDL (Zhang et al., 2020; Legried et al., 2020; Markin and Eulenstein, 2020), while FastMulRFS has been proved to be statistically consistent under a model of either only duplication or only loss (Molloy and Warnow, 2020).

Given a collection of trees corresponding to gene families (one tree per gene family), we generated four types of input to each of the methods:

- ONLY: The input consists of trees of *only* gene families that are present in exactly one copy in each of the species.
- ORTHOLOGS: The input consists of trees of ONLY gene families that have no history of gene duplication. These are canonical single-copy orthologs.
- ONE: The input consists of trees of *all* gene families, but where a single copy per species per gene family is selected at random and the remaining copies are removed. If a gene family has no copies at all for some species, then the resulting tree of that gene family also has no copies for that species.
- ALL: The input consists of trees of *all* gene families, but where all copies of a gene in a species are treated as multiple alleles from different individuals within the species. Similar to ONE, if a gene family has no copies at all for some species, then the resulting tree of that gene family also has no copies for that species.

ONLY corresponds to the practice that is followed in many phylogenomic studies, though it does not necessarily guarantee that the included genes are orthologs. Instead, “hidden paralogs” (Doolittle and Brown, 1994) or “pseudoorthologs” (Koonin, 2005) may occur: these are cases in which complementary losses result in single-copy paralogs present in different species. ORTHOLOGS corresponds to a scenario where researchers know which genes have a history of duplication and can exclude them from their analysis. ONE is likely to have some hidden paralogs in the input, unless GDL does not occur. By construction, ALL has all orthologs and paralogs as input, but these are effectively labeled as orthologs, with the (wrong) implied assumption that multiple individuals are sampled per species.

### Simulation setup

For model species trees, we used the trees of 16 fungal species and 12 fly species reported in Rasmussen and Kellis (2012) and shown in Fig. 1. The 16 fungal species are:*Candida albicans* (Calb), *Candida tropicalis* (Ctro), *Candida parapsilosis* (Cpar), *Lodderomyces elongisporus* (Lelo), *Candida guilliermondii* (Cgui), *Debaryomyces hansenii* (Dhan), *Candida lusitaniae* (Clus), *Saccharomyces cerevisiae* (Scer), *Saccharomyces paradoxus* (Spar), *Saccharomyces mikatae* (Smik), *Saccharomyces bayanus* (Sbay), *Candida glabrata* (Cgla), *Saccharomyces castellii* (Scas), *Kluyveromyces lactis* (Klac), *Ashbya gossypii* (Agos), and *Kluyveromyces waltii* (Kwal). Note that *Saccharomyces castellii* has since been re-named *Naumovozyma castellii* (https://www.uniprot.org/taxonomy/27288), *Kluyveromyces waltii* has since been re-named *Lachancea waltii* (https://www.uniprot.org/taxonomy/1089441), and *Ashbya gossypii* has been re-named *Eremothecium gossypii* (https://www.uniprot.org/taxonomy/33169).

**Fig. 1.**
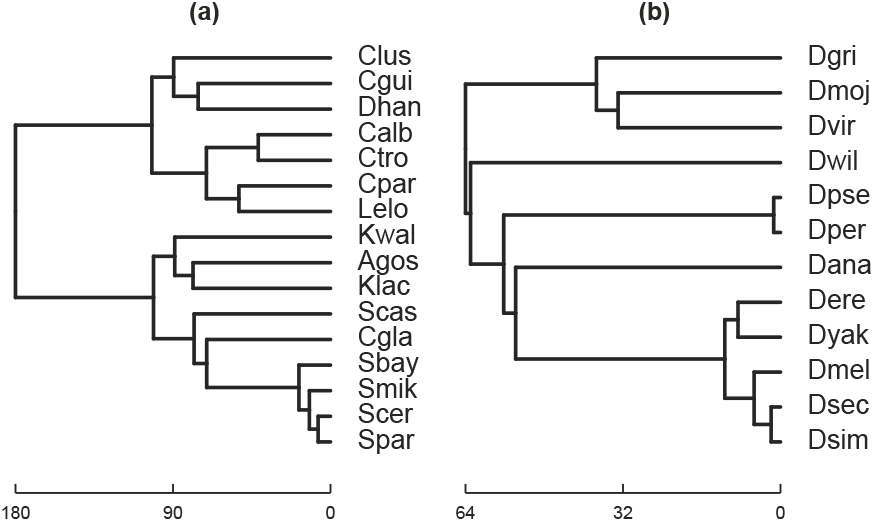
The species trees reported in Rasmussen and Kellis (2012), which we use as the topologies in the simulations and in the empirical data analysis. (a) The species tree of 16 fungal species. (b) The species tree of 12 fly species. The species tree topologies and their branch lengths in units of million years are taken from http://compbio.mit.edu/dlcoal/.

The 12 fly species are: *Drosophila melanogaster* (Dmel), *Drosophila simulans* (Dsim), *Drosophila sechellia* (Dsec), *Drosophila erecta* (Dere), *Drosophila yakuba* (Dyak), *Drosophila ananassae* (Dana), *Drosophila pseudoobscura* (Dpse), *Drosophila persimilis* (Dper), *Drosophila willistoni* (Dwil), *Drosophila mojavensis* (Dmoj), *Drosophila virilis* (Dvir), and *Drosophila grimshawi* (Dgri).

To generate gene trees while allowing for ILS and GDL, we used SimPhy (Mallo et al., 2015) with the parameters specified below (assuming all species are diploid). SimPhy uses the three-tree model developed in Rasmussen and Kellis (2012) to simulate data. In this model, a *locus tree* is simulated within the branches of the species tree. All incongruence between the locus tree and the species tree is due to GDL. Then, a *gene tree* is simulated within the branches of the locus tree, where all incongruence between the locus tree and the gene tree is due to ILS. The resulting gene tree differs from the species tree due to a combination of ILS and GDL. Using the locus trees as input to an inference method amounts to using data where all incongruence is solely due to GDL (but not ILS). Setting the rates of GDL to 0 amounts to generating gene trees where all incongruence is solely due to ILS. Note that SimPhy makes two further assumptions relevant to the results presented here: first, it assumes no hemiplasy of new duplication mutations. That is, all new duplicates immediately fix before they can be lost during a polymorphic phase. Rasmussen and Kellis (2012) found that this assumption affected 5% of gene families simulated under similar conditions. Furthermore, hemiplasy results in an excess of apparent gene losses, which should not affect inferences of species trees. The second assumption is that all gene families are independent: no events duplicate or delete more than a single gene at a time. In real data, large-scale events (including whole-genome duplications) can affect many genes at a time.

For the fungal tree simulated datasets, we used five different duplication and loss rates (assuming duplication and loss rates are equal): 0 (to investigate the performance when ILS, but not GDL, acted on the gene families), 1 × 10^−10^, 2 × 10^−10^, 5 × 10^−10^, and 10 × 10^−10^ per generation. We take the case where the rate is 1 × 10^−10^ to be similar similar to the duplication rate of 7.32 × 10^−11^ and loss rate of 8.59 × 10^−11^ used by Rasmussen and Kellis (2011), and denote this rate as “1x”. We used two effective population sizes: 10^7^ and 5 × 10^7^, where the former was also used by Rasmussen and Kellis (2012) as the true population size. We assumed 0.9 years per generation as in Rasmussen and Kellis (2012) and used 4 × 10^−10^ as the nucleotide mutation rate per site per generation, similar to the rates of 3.3 × 10^−10^ and 3.8 × 10^−10^ used by Zhang and Wu (2017) and Lang and Murray (2008), respectively.

For the fly tree simulated datasets, we used five different duplication and loss rates (assuming duplication and loss rates are equal): 0, 1 × 10^−10^, 2 × 10^−10^, 5 × 10^−10^, and 10 × 10^−10^ per generation. A GDL rate of 1.2 × 10^−10^ was used in (Rasmussen and Kellis, 2012; Zhang and Wu, 2017) and reported by Hahn et al. (2007); we again denote this rate as “1x”. We used two effective population sizes: 10^6^ and 5 × 10^6^, similar to the values used in (Rasmussen and Kellis, 2012) and the estimated value of 1.15 × 10^6^ reported in (Sawyer and Hartl, 1992; Pollard et al., 2006). We assumed 10 generations per year as in (Rasmussen and Kellis, 2012; Zhang and Wu, 2017) and used 3 × 10^−9^ as the mutation rate per site per generation, similar to the rate of 5 × 10^−9^ found in Schrider et al. (2013).

For each combination of GDL rate and population size, 10,000 gene families (each containing a locus tree and its corresponding gene tree) were simulated in this fashion as one dataset. Ten such data sets, each with 10,000 gene families, were generated for each condition. To study the effect of using datasets of varying sizes, for each of the 10 datasets we randomly sampled 10, 50, 100, and 250 gene families from the 10,000 gene families under the ALL, ONE, ONLY, and ORTHOLOGS scenarios. In case the number of available gene families that fits ONLY or ORTHOLOGS is smaller than the desired size, that number of gene families was used (e.g., when only 6 gene family trees are available when data sets of size 10 are desired, the 6 trees are used as input).

To study the effect of GDL and ILS on species tree estimates, for each dataset of trees (true gene trees or true locus trees; that is, trees without estimation error) of a given size, we fed the dataset as input to `InferNetwork_MPL`, ASTRAL, NJ_st_, ASTRAL-Pro, and FastMulRFS and computed the Robinson-Foulds distance (Robinson and Foulds, 1981), normalized by the number of internal branches in the (unrooted) species tree to obtain a value between 0 and 1. This is the normalized distance between the true and inferred species trees. To study the further effect of error in the gene tree estimates on species tree estimates, we simulated the evolution of sequences of length 500 nucleotides on all gene trees under the HKY model, using Seq-gen (Rambaut and Grassly, 1997). We then inferred gene trees from the simulated sequence data using IQ-TREE (Nguyen et al., 2014). Furthermore, to study the effect of error in the locus tree estimates, we treated the true locus tree as a gene tree and simulated the evolution of sequences of length 500 nucleotides on all locus trees under the HKY model, again using Seq-gen, and inferred locus trees from the simulated sequence data using IQ-TREE. It is important to note that in practice only gene trees, but not locus trees, are inferrable, as the locus tree is an artifact of the three-tree model and not a biological entity (Rasmussen and Kellis, 2012). However, conducting analysis using inferred locus trees gives a picture of the performance when all incongruence is due to GDL and gene tree error only. Finally, `InferNetwork_MPL` assumes that the input gene trees are rooted. In this study, we rooted the gene tree estimates by minimizing deep coalescences (Maddison, 1997; Than and Nakhleh, 2009); that is, we rooted each gene tree in a way that minimizes the number of extra lineages when reconciled with the true species tree.

### Biological data

For the fungal dataset, we used 2932 gene trees reported in http://compbio.mit.edu/dlcoal/ and estimated with PhyML (Guindon and Gascuel, 2003), where 1867 gene trees fit the ONLY setting. For the fly dataset, we used 9233 gene trees from (Hahn et al., 2007) reconstructed using the neighbor-joining algorithm, where 6698 gene trees fit the ONLY setting. For the fly dataset, we removed any gene trees containing polytomies prior to running NJst. In neither dataset did we attempt to identify single-copy orthologs. We again rooted each gene tree in the empirical data with respect to the species trees of Fig. 1 so as to minimize deep coalescences (Maddison, 1997; Than and Nakhleh, 2009) using the method of Yu et al. (2011), as implemented by the function ProcessGT in PhyloNet (Wen et al., 2018). We estimated species trees using ASTRAL, NJ_st_, maximum pseudo-likelihood, ASTRAL-Pro, and FastMulRFS with these gene trees as input.

## RESULTS

### Characteristics of the simulated data

Before we describe the inference results, we discuss the characteristics of the simulated data. First, we investigated the effects of gene duplication and loss on the number of gene copies per species in each gene family. Fig. 2(a-b) and fig. S1(a-b) show data on the sizes (numbers of copies) of gene families in the 16-taxon and 12-taxon data sets, respectively, under the various settings of effective population sizes and duplication and loss rates.

**Fig. 2.**
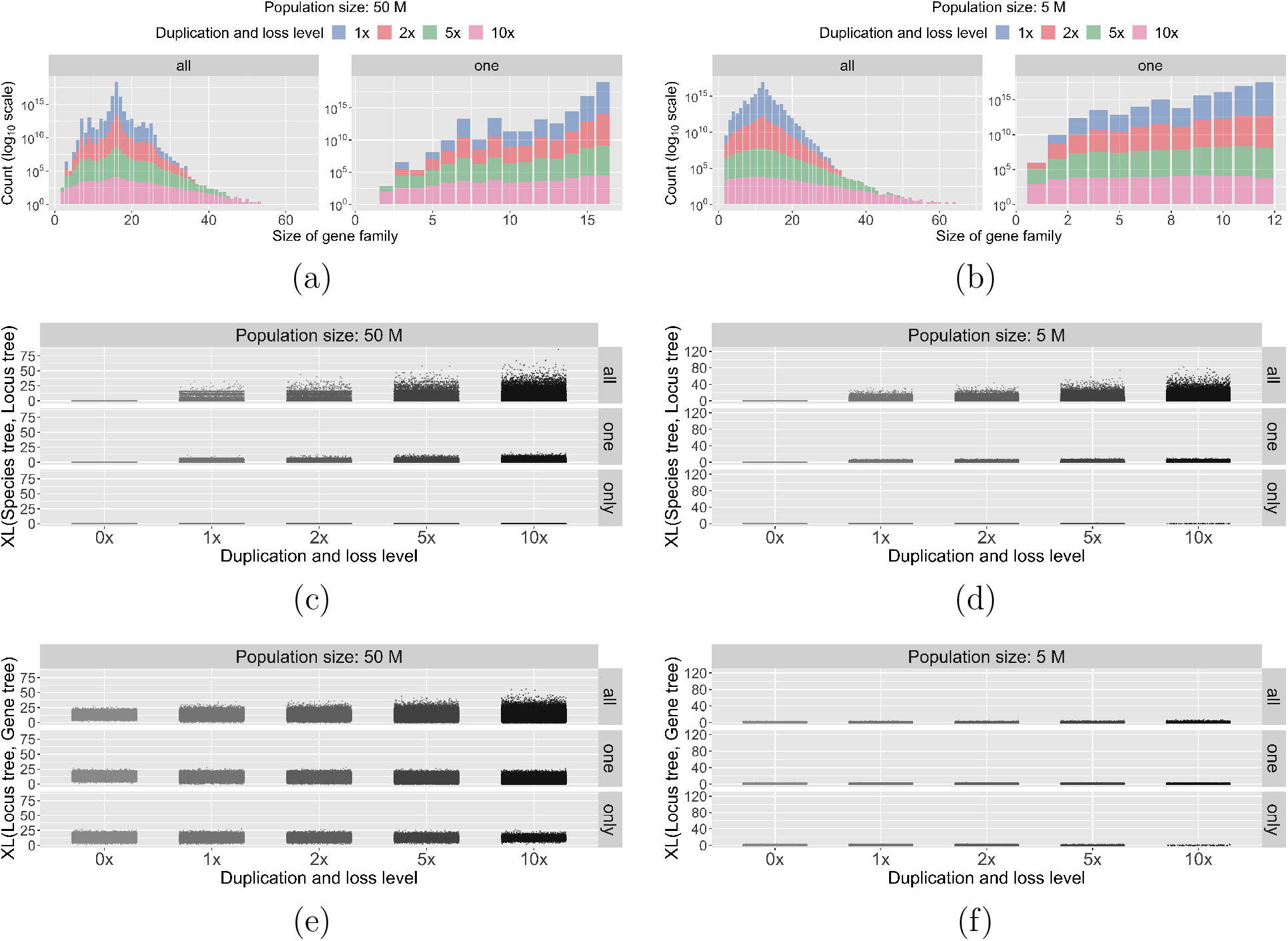
Characteristics of the simulated data under different settings of the duplication/loss rates and tree topologies. The duplication/loss rates are denoted by the rate multiplier (0x, 1x, 2x, 5x and 10x), where 1x is the rate found in nature for the clade represented by each species tree topology (see Methods). (a-b) Distribution of the total number of gene copies in individual gene families in the 16-taxon and 12-taxon data sets, respectively. Note that the two tree topologies also have different simulated effective population sizes in these figures (see fig. S1(a-b) for more conditions). (c-d) Scatter plots of XL(Species tree, Locus tree), the number of extra lineages when reconciling the true locus trees with the true species tree, for the 16-taxon and 12-taxon data sets, respectively. These plots therefore represent the effects of GDL alone. (e-f) Scatter plots of XL(Locus tree, Gene tree), the number of extra lineages when reconciling the true gene trees with the true locus tree, for the 16-taxon and 12-taxon data sets, respectively. These plots therefore represent the effects of ILS alone, though note that higher rates of GDL allow there to be more gene tree branches on which ILS can act.

Clearly, the higher the GDL rates, the larger the variance in size of gene families. The figure also shows that the average size of a gene family is roughly equal to the number of species, with the largest gene families having 65 copies for the 16-taxon datasets, and 94 copies for the 12-taxon datasets (recall that these trees use different rates of GDL). We then counted the average (over the 10 datasets per setting) number of gene families for each setting that have ONLY one copy per species and the average number of gene families with no history of duplication (i.e. ORTHOLOGS). The results are shown in Table 1. The table shows that as the GDL rates increase, the number of single-copy orthologs decreases. However, as predicted by theory (Smith and Hahn, 2021a), there appear to be very few pseudoorthologs in the ONLY dataset.

**Table 1.** The average number of gene families that fit the ONLY/ORTHOLOGS settings out of the 10,000 gene families.

| GDL rate | $N_e$ | 16-taxon data | | 12-taxon data | |
| --- | --- | --- | --- | --- | --- |
| | | $10^7$ | $5 \times 10^7$ | $10^6$ | $5 \times 10^6$ |
| $1 \times 10^{-10}$ | | 7619/7616 | 7585/7583 | 4591/4554 | 4584/4550 |
| $2 \times 10^{-10}$ | | 5794/5782 | 5787/5775 | 2197/2131 | 2176/2111 |
| $5 \times 10^{-10}$ | | 2554/2521 | 2538/2508 | 268/226 | 266/222 |
| $1 \times 10^{-9}$ | | 689/659 | 688/657 | 12/6 | 13/7 |

We then set out to assess the extent of incongruence in the gene trees due to GDL and ILS. For every pair of true species tree and true locus tree, we computed the number of extra lineages (Maddison, 1997) using the DeepCoalCount_tree command in PhyloNet (Than and Nakhleh, 2009; Wen et al., 2018) as a proxy for the amount of incongruence in the data. Here, we treated all gene copies from the same species as different individuals. Zero extra lineages mean there is no incongruence between the two trees, and the higher the value, the more incongruence there is. In particular, no incongruence means that all gene copies from the same species are monophyletic in the locus tree, and when restricted to a single arbitrary copy per species, the locus tree and species tree have identical topologies.

Fig. 2(c-d) and fig. S1(c-d) show data on the number of extra lineages in the simulated 16-taxon and 12-taxon datasets, respectively, under the various settings of effective population sizes and duplication and loss rates. It is important to note that all incongruence in this case is exclusively due to GDL (ILS is not a factor in the results in these two panels). The panels do not have results for the GDL rate of 0x, because in such cases there is no incongruence at all between the locus tree and the species tree, and thus there are zero extra lineages. The results show that, unsurprisingly, there is much more incongruence for the ALL scenario than the ONE scenario. For the ONLY scenario, there is very little incongruence in either dataset.The incongruence in ONLY would indicate the phenomenon of hidden paralogy: single-copy genes are paralogs, so that their gene trees do not always agree with the species tree. Given the small number of hidden paralogs (Table 1), these results are unsurprising. The ORTHOLOGS datasets are not plotted, because the number of extra lineages in those locus trees is always zero, as expected.

We also computed the number of extra lineages when reconciling the true gene trees with the true locus trees. Here, incongruence is exclusively due to ILS (GDL is not a factor). Fig. 2(e-f) and fig. S1(e-f) show data on the number of extra lineages in the simulated 16-taxon and 12-taxon datasets, respectively, under the various settings of effective population sizes and duplication and loss rates. When the gene tree topology is identical to the locus tree topology, the number of extra lineages is zero, and the larger the number of extra lineages, the more ILS has an effect on the data. The figure shows that, as expected, the amount of ILS is larger for larger population sizes, and consequently there is much more ILS in the 16-taxon dataset than in the 12-taxon dataset. One other trend to observe is that, on average, the amount of incongruence due to ILS increases with the increase in the GDL rate. This is a reflection of the fact that for higher GDL rates, the locus trees are larger (more leaves and internal branches) and this naturally results in more branches that can be affected by ILS. Finally, the amount of incongruence due to ILS is generally far lower than the amount due to GDL in the 12-taxon dataset, while the levels of incongruence due to GDL and ILS are similar in the 16-taxon dataset, especially when the rates of duplication and loss are high.

### Results on Simulated Data

We are now in position to describe the inference results. We show figures for the 16-taxon datasets in the main text, while figures for the 12-taxon datasets are all in the Supplementary Materials (figs. S8 to S11). The results for the 12-taxon datasets are consistently better in terms of accuracy, so we chose to focus here on the less-optimal results.

We first ran the inference methods ASTRAL, `InferNetwork_MPL`, NJ_st_, ASTRAL-Pro, and FastMulRFS on the true gene trees for all four input scenarios: ALL, ONE, ONLY, and ORTHOLOGS. In this case, gene tree estimation error is not a cause of gene tree incongruence. Instead, all incongruence is due to a combination of ILS and GDL. Results on the full 16-taxon tree are shown in Fig. 3 and fig. S4. Note that, in all cases, using input data with GDL levels of 0 amounts to inferring a species tree from gene trees whose incongruence is solely due to ILS.

**Fig. 3.**
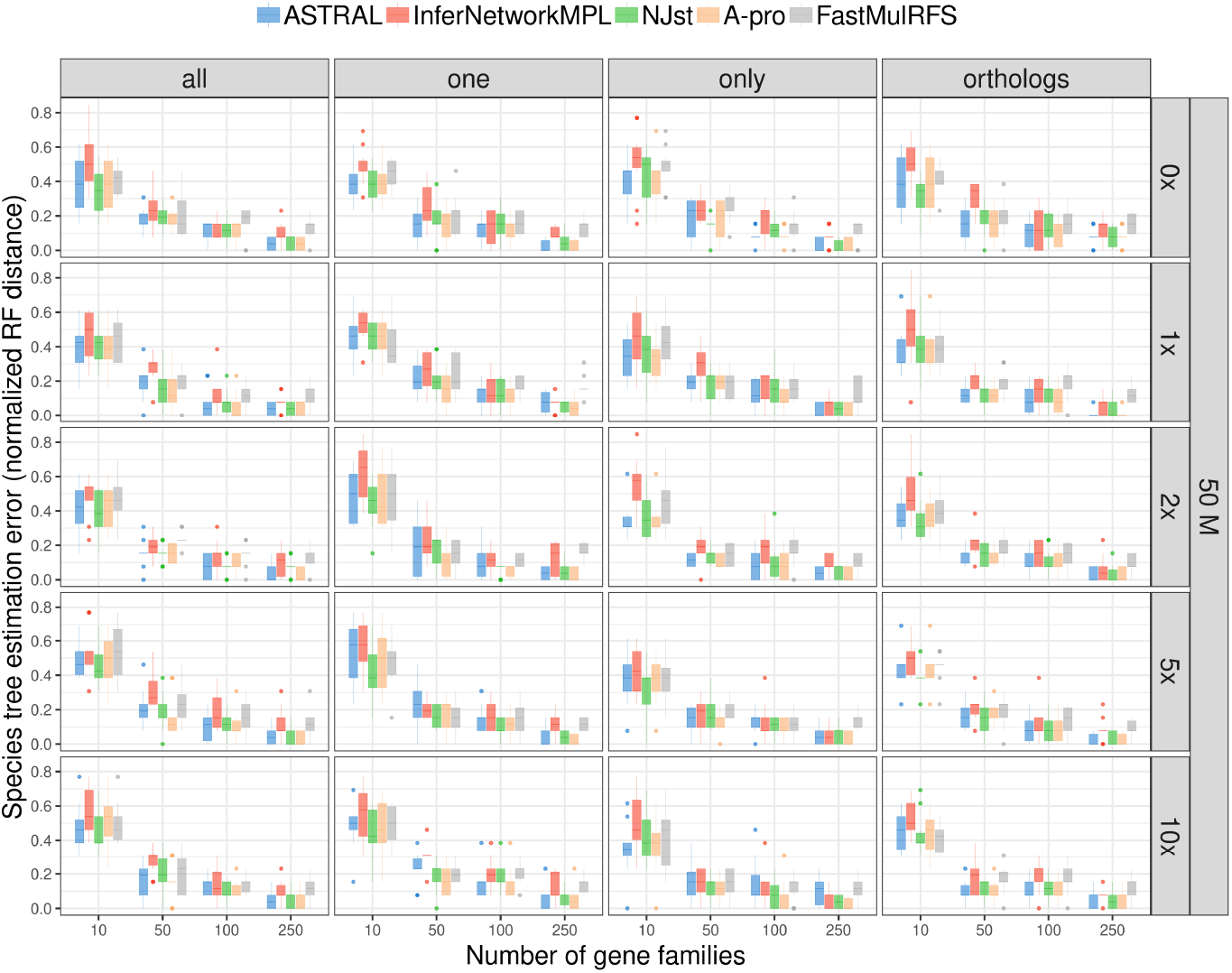
The normalized RF distances between the true species tree and the ones inferred from true gene trees. These simulations therefore include the effects of both ILS and GDL (but no gene tree estimation error). The five inference methods used are ASTRAL, `InferNetwork_MPL`, NJ_st_, ASTRAL-Pro (“A-pro”), and FastMulRFS), and data is simulated on the 16-taxon tree with a population size of 5.0 × 10^7^ and varying GDL rates. The duplication/loss rates are denoted by the rate multiplier (0x, 1x, 2x, 5x and 10x), where 1x is the rate estimated in nature for fungi. Each row corresponds to a combination of population size and GDL rates. The X-axis in each panel represents the number of gene families used and the Y-axis represents the normalized RF distance.

There are several observations based on these results. First, the accuracy of the inferred 16-taxon trees is much lower in general than that of the inferred 12-taxon trees. In particular, for the 12-taxon data sets, the species trees are perfectly estimated in almost all cases (fig. S3), whereas the species tree estimation error is high, especially for the larger population sizes, for the 16-taxon data sets. As shown in Fig. 2 and fig. S1, both datasets have similar gene family sizes, but differ significantly in terms of the amount of ILS in the data, with the 12-taxon datasets having very little ILS. Therefore, the straightforward explanation for the observed differences species tree inference accuracy between the 16- and 12-taxon data sets is the higher level of ILS in the former. Given that the level of incongruence due to GDL is similar between the 16-taxon and 12-taxon data sets (Fig. 2(c-d) and fig. S1(c-d)), these results point to the larger role that ILS plays in the methods’ performance than GDL does.

Second, in the case of the 16-taxon data, the performance of all methods improves as the number of gene families used as input to the method increases. Note also that the largest dataset used here consists of only 250 gene trees, which is much smaller than the number available in most phylogenomic data sets. While there is very little difference observed in the performance among the methods on the 16-taxon data, ASTRAL, ASTRAL-Pro, and NJ_st_ are more similar to each other in terms of performance than either of them is to inference under maximum pseudo-likelihood or FastMulRFS. This makes sense as ASTRAL, ASTRAL-Pro, and NJ_st_ are summary methods that make inference based on statistics derived from the input gene trees, whereas maximum pseudo-likelihood uses calculations based on the multispecies coalescent directly. The performance of FastMulRFS is similar to that of other methods, but its error rates remain higher than the other methods when more gene families are used. Although ASTRAL-Pro and FastMulRFS were developed with gene duplication and loss in mind, they do not appear to outperform the other summary methods.

Third, the level of ILS for a population size of 50M is higher than for a population size of 10M, and this results in lower accuracy of inferred species trees by all methods in the former case (fig. S4). This behavior is expected for any method, regardless of whether GDL is acting. Notably, FastMulRFS was not developed to deal correctly with ILS, and seems to have an inflated error rate with larger population sizes, but not with smaller population sizes (fig. S4), suggesting that ILS may be the cause of higher error rates in this method.

Lastly, we observe very little difference in the accuracy of inferred species trees across the four input scenarios: ALL, ONE, ONLY, and ORTHOLOGS. The only case in which there is a noticeable difference is in the 12-taxon datasets with the duplication rate 10x that found in nature, when only ten genes are used for inference (figs. S8 and S9). These results imply that the presence of paralogs in the data, no matter how they are treated, does not have much of an effect on the performance of the five methods, unless very few genes are used.

The results thus far raise the important question: Does GDL have any effect on the performance of these five methods? To answer this question, we ran all of them on the locus trees as input to infer species trees. By the three-tree model, this amounts to feeding these methods “gene trees” whose incongruence is solely due to GDL; that is, ILS plays no role in incongruence here. It is important to point out that locus trees are mathematical constructs of the three-tree model; in practice, inferring a locus tree is not possible, unless the data has no ILS at all. We conducted this experiment to study the performance of methods when GDL, but not ILS, causes all incongruence. Results on the full 16-taxon datasets are shown in Fig. 4 and fig. S5. As the results show, all methods infer the species tree perfectly accurately on almost all data sets, regardless of the parameter settings and the input scenario. In other words, when these methods—some of which have been developed based on the multispecies coalescent directly (`InferNetwork_MPL`), some of which were inspired by the MSC (ASTRAL, ASTRAL-Pro, and NJ_st_), and one that does not deal with ILS at all (FastMulRFS)—are applied to data that have no ILS but do have paralogs in them, they have almost perfect accuracy in terms of the species tree topology they infer, under the conditions of our simulations. Combined with the results summarized in Fig. 3 and fig. S4, these results show, perhaps surprisingly, that methods developed to handle ILS but not GDL do much better in handling GDL than they do in handling ILS. Perhaps unsurprisingly, ASTRAL-Pro and FastMulRFS, methods designed to handle GDL, also perform well on the true locus trees. The inflated errors seen with FastMulRFS under some settings with gene trees are absent when true locus trees are used as input, suggesting that, indeed, these errors were due to ILS. ASTRAL-Pro was designed to deal with both ILS and GDL and performs well on both true gene trees and true locus trees.

**Fig. 4.**
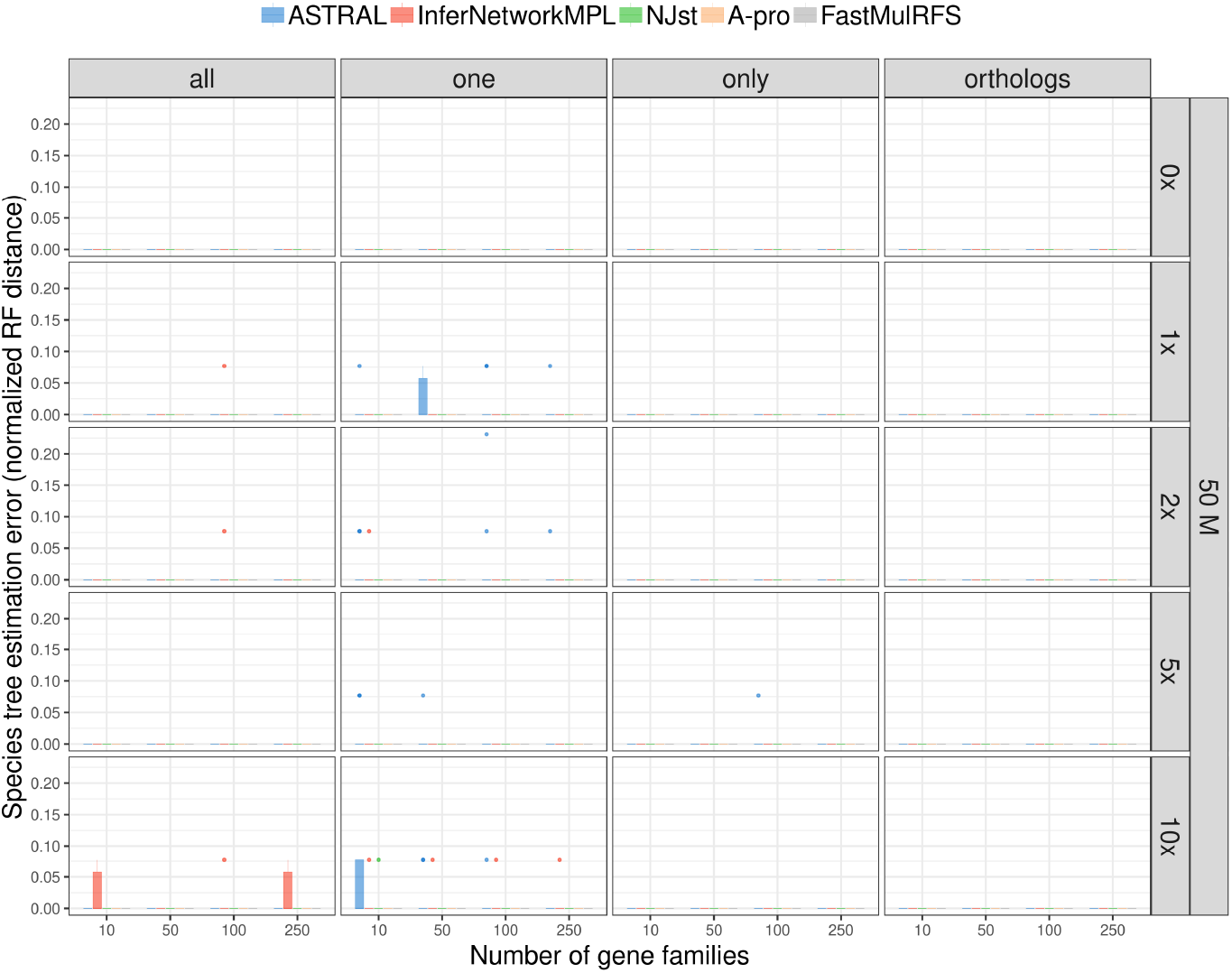
The normalized RF distances between the true species tree and the ones inferred from true locus trees. These simulations therefore include the effects of GDL only (no ILS or gene tree estimation error). The five inference methods used are ASTRAL, `InferNetwork_MPL`, NJ_st_, ASTRAL-Pro (“A-pro”), and FastMulRFS), and data is simulated on the 16-taxon tree with a population size of 5.0 × 10^7^ and varying GDL rates. The duplication/loss rates are denoted by the rate multiplier (0x, 1x, 2x, 5x and 10x), where 1x is the rate estimated in nature for fungi. Each row corresponds to a combination of population size and GDL rates. The X-axis in each panel represents the number of gene families used and the Y-axis represents the normalized RF distance.

In practice, gene trees are unknown and are inferred from sequence data. Therefore, to simulate more realistic scenarios, we inferred gene trees and locus trees from simulated sequence data and fed these tree estimates as input to the five methods. In this case, gene tree estimation error is a factor in the observed incongruences. We show the extent of error in the estimated gene and locus trees for the 16-taxon data in fig. S2. Gene tree estimation error is measured by the normalized RF distance between the true gene tree and the reconstructed gene tree. For the 12-taxon data set, the average gene tree estimation error ranges from 0.456 to 0.648, whereas the average locus tree estimation error is slightly lower, ranging from 0.414 to 0.627 (fig. S3). For the 16-taxon data set, the average gene tree estimation error ranges between 0.073 to 0.130 while the average locus tree estimation error ranges from 0.065 to 0.099. In other words, there is much less gene tree estimation error in the 16-taxon data sets than in the 12-taxon data sets.

Results of species tree inference using the full 16-taxon dataset based on estimated gene trees are shown in Fig. 5 and fig. S6; those based on the locus tree estimates are shown in Fig. 6 and fig. S7. These results should be contrasted with Fig. 3, fig. S4, Fig. 4 and fig. S5, respectively, to understand the effect of gene tree estimation error on the accuracy of species tree inference.

**Fig. 5.**
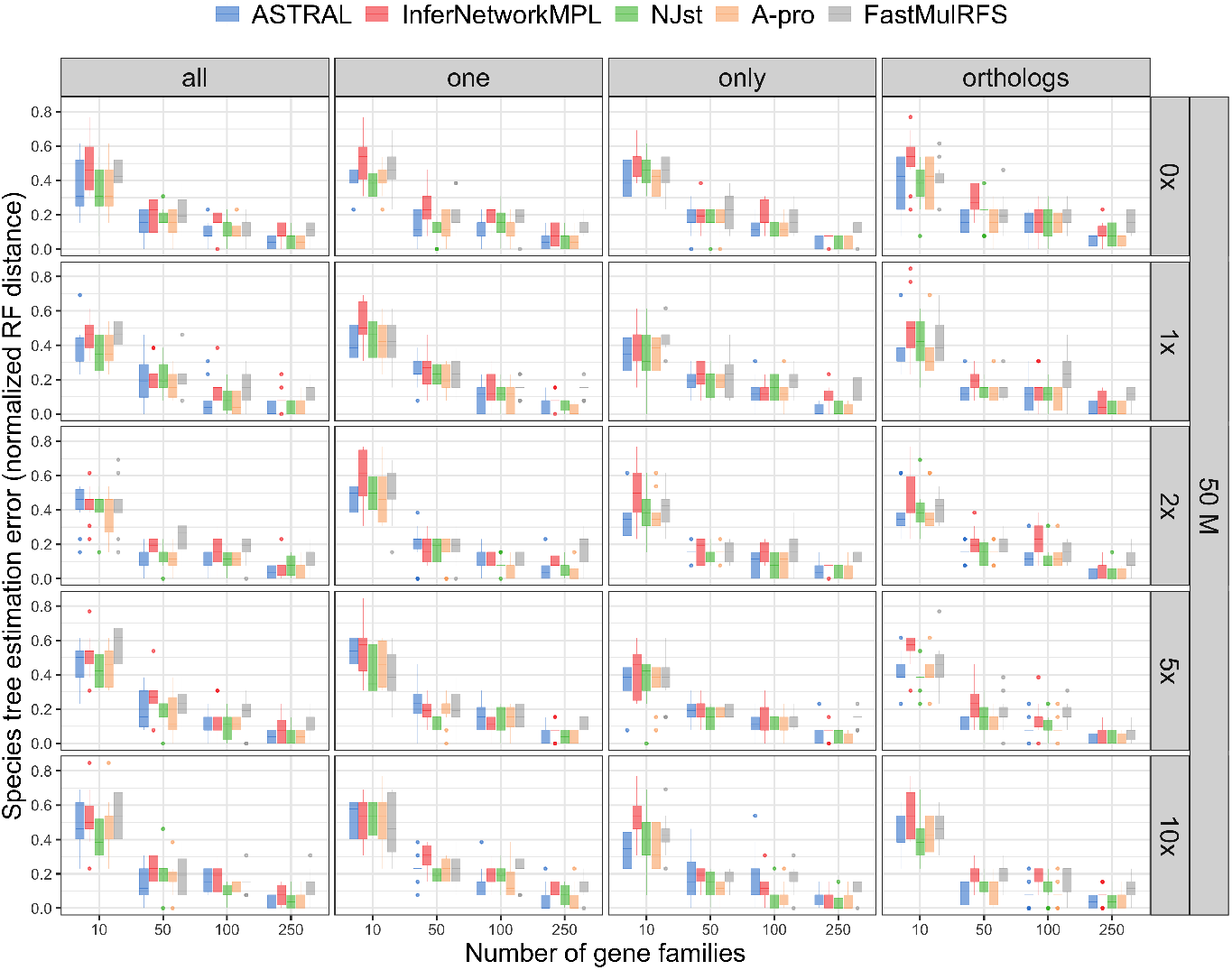
The normalized RF distances between the true species tree and the ones inferred from estimated gene trees. These simulations therefore include the effects of GDL, ILS, and gene tree estimation error. The five inference methods used are ASTRAL, `InferNetwork_MPL`, NJ_s_t, ASTRAL-Pro (“A-pro”), and FastMulRFS), and data is simulated on the 16-taxon tree with a population size of 5.0 × 10^7^ and varying GDL rates. The duplication/loss rates are denoted by the rate multiplier (0x, 1x, 2x, 5x and 10x), where 1x is the rate estimated in nature for fungi. Each row corresponds to a combination of population size and GDL rates. The X-axis in each panel represents the number of gene families used and the Y-axis represents the normalized RF distance.

**Fig. 6.**
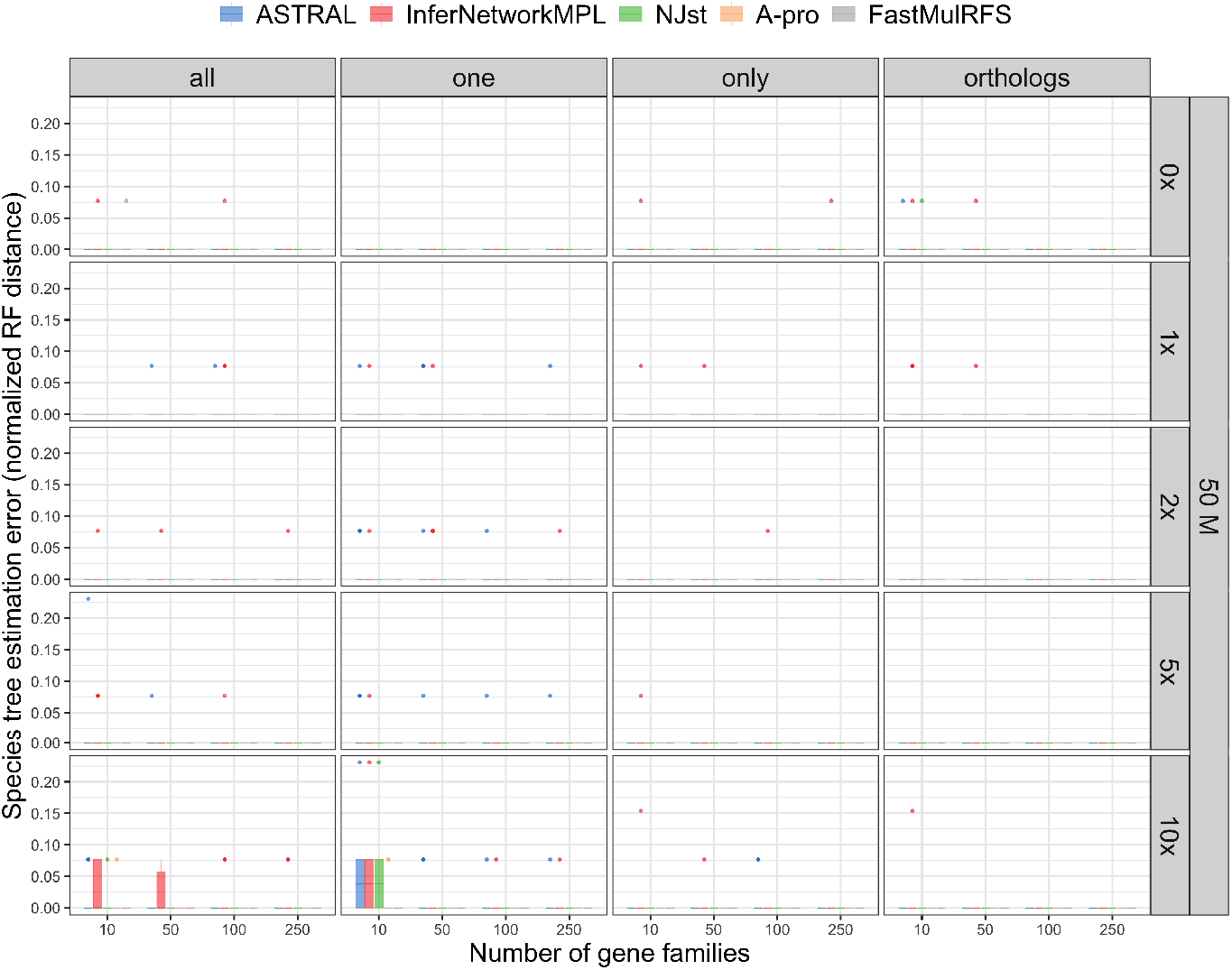
The normalized RF distances between the true species tree and the ones inferred from the estimated locus trees. These simulations therefore include the effects of GDL and gene tree estimation error (no ILS). The five inference methods used are ASTRAL, `InferNetwork_MPL`, NJ_st_, ASTRAL-Pro (“A-pro”), and FastMulRFS), and data is simulated on the 16-taxon tree with a population size of 5.0 × 10^7^ and varying GDL rates. The duplication/loss rates are denoted by the rate multiplier (0x, 1x, 2x, 5x and 10x), where 1x is the rate estimated in nature for fungi. Each row corresponds to a combination of population size and GDL rates. The X-axis in each panel represents the number of gene families used and the Y-axis represents the normalized RF distance.

In the case of species tree inferences using data where ILS, GDL, and gene tree estimation error are involved, the error rates of all five species tree inference methods went up, as expected (Fig. 5 and fig. S6), but only slightly. The accuracy of the species trees improves as the number of gene families increases. As discussed above, the error in gene tree estimates in the 16-taxon datasets is very low. Since gene tree estimation error in the 12-taxon datasets is much higher (because the higher substitution rates result in noisier sequence data), we observe a larger impact of this error on the performance of methods on the 12-taxon datasets (fig. S10). While the methods had an almost perfect accuracy on true gene trees, species tree estimates now have as high as 50% error when 10 gene family trees are used, and close to 25% error when 250 gene family trees are used (fig. S10). These results illustrate the large impact gene tree estimation error has on these methods. In the case of the 12-taxon datasets, the impact of gene tree estimation error significantly outweighs that of ILS or GDL.

Fig. 6 and fig. S7 demonstrate how GDL and gene tree estimation error (but no ILS) impact species tree inference. As with Fig. 4 and fig. S5, which show almost perfect performance of species tree inference from true locus trees (i.e., GDL and no ILS), we observe little reduction in performance here due to error in the estimates of gene trees. The results demonstrate that in the absence of ILS, all methods are robust to gene tree estimation error, except when the number of gene families is very small. In the case of the 12-taxon datasets, where locus tree estimation error is much higher, the five species tree inference methods also have comparable, but lower, accuracies (fig. S11).

All of these results combined point to a very small impact of GDL on the performance of the five studied species tree inference methods and given the simulation parameters used here, regardless of how the paralogs are handled. On the other hand, across all datasets it was evident that gene tree estimation error has a noticeable impact on the methods’ performance, and that ILS often had a substantial impact on accuracy.

### Results on Biological Data

We ran all five methods used above on two empirical datasets, each consisting of thousands of gene trees. As the two datasets were the basis for the simulated data presented above, we believe that they share many of the same properties as these data.

For the 16 fungal genomes, the inferred species trees from all five methods differ from the tree shown in Fig. 1(a). ASTRAL, NJ_st_, ASTRAL-Pro and FastMulRFS inferred the same topology depicted in Fig. 7(c) under all three input scenarios (recall that ORTHOLOGS is not used here, since true orthologs are not known). The same phylogeny is also inferred by `InferNetwork_MPL`(ONE). This inferred tree is topologically different from the tree shown in Fig. 1(a): in particular, the positions of *Kluyveromyces waltii* and *Kluyveromyces lactis* have been switched, as have the positions of *Candida glabrata* and *Saccharomyces castellii* (Fig. 7(c)). The trees inferred by `InferNetwork_MPL`(ALL) and `InferNetwork_MPL`(ONLY) differ from the reference tree of Fig. 1(a) in terms of the placement of *Candida glabrata* and *Saccharomyces castellii*, as shown in Fig. 7(a) and Fig. 7(b). `InferNetwork_MPL`(ALL) additionally grouped *Saccharomyces cerevisiae* and *Saccharomyces mikatae* as sisters, and switched the position of *Kluyveromyces waltii* and *Kluyveromyces lactis.* Interestingly, the position of *Candida glabrata* is not a settled issue: Shen et al. (2016) label the relevant branch as “unresolved” in their analysis of 1,233 single-copy orthologs. Similarly, their results support the same placement of *Kluyveromyces lactis* as in Figs 7a and 7c here. The placement of these species shown in Fig. 1(a) originally comes from a concatenated analysis of 706 single-copy genes (Butler et al., 2009).

**Fig. 7.**
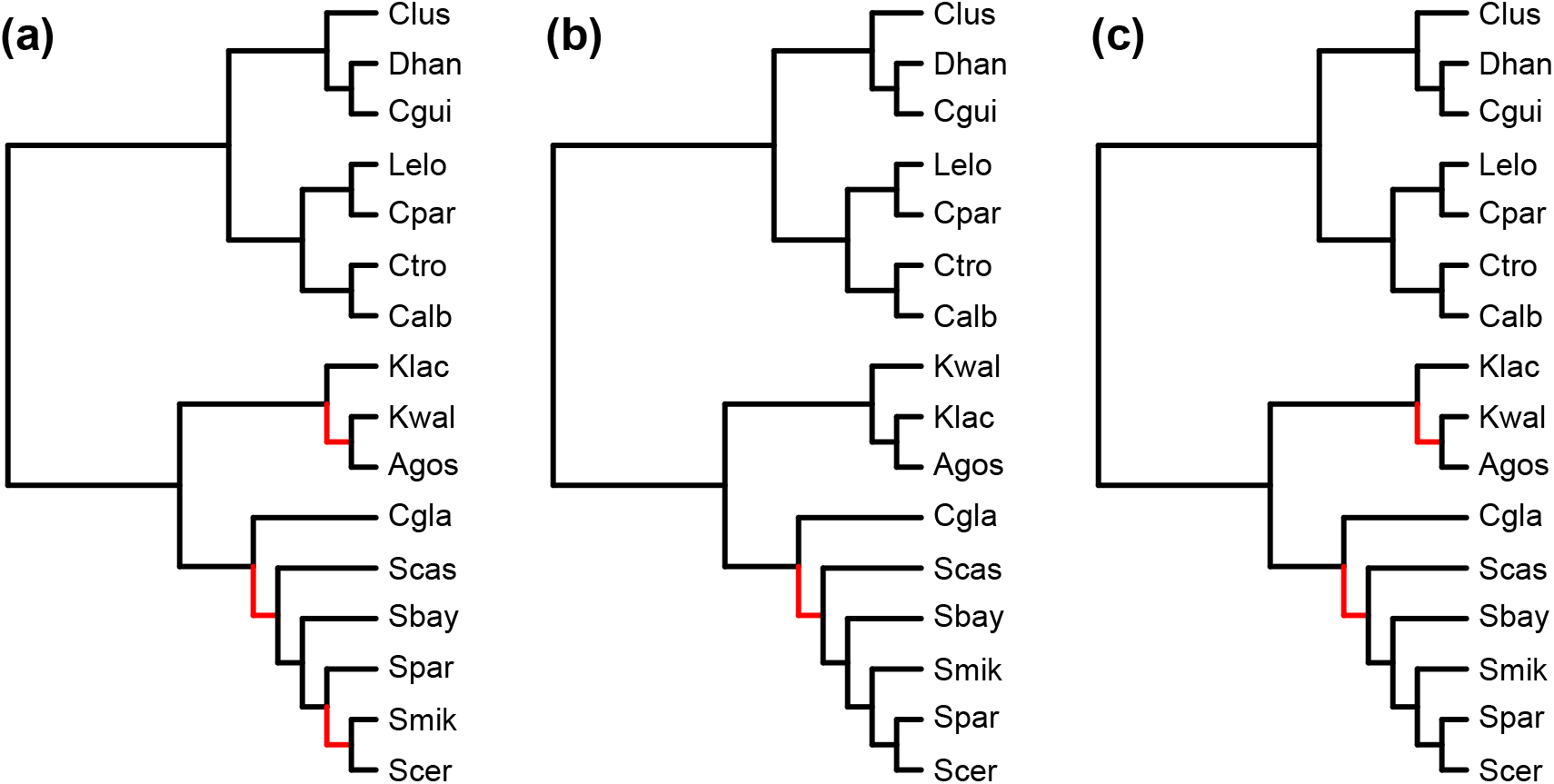
Inferred fungal species trees. (a) The fungal species tree inferred by `InferNetwork_MPL`(ALL). (b) The fungal species tree inferred by `InferNetwork_MPL`(ONLY) (c) The fungal species tree inferred by ASTRAL, NJ_st_, ASTRAL-Pro, FastMulRFS, and `InferNetwork_MPL`(ONE). Differences between the inferred species trees and the tree in Fig. 1 are highlighted in red.

For the 12 fly genomes, all three sampling schemes and all five methods inferred the exact same tree as the species tree shown in Fig. 1(b).

## Discussion

As phylogenomic datasets grow, our ability to use them within the bounds of current analysis paradigms shrinks. One of the main problems is the decreasing number of gene families that are single-copy as the number of sampled species increases (Emms and Kelly, 2018). Because most current phylogenetic methods assume that only single-copy orthologs are being used, this restriction means that such methods cannot be used for datasets with even several dozen taxa without severe downsampling or other *ad hoc* solutions (e.g., Thomas et al., 2020). Here, we set out to ask whether phylogenomic methods intended to deal with incongruence due to ILS can be applied to data containing both orthologs and paralogs, which contain incongruence due to GDL.

On simulated datasets where only ILS acted, and GDL was not a factor, all methods had the expected performance: accurate species tree estimates that improved as the number of gene trees used increases. In the case where the level of ILS was very low (the 12-taxon data), the methods had perfect performance under almost all conditions, regardless of the number of gene trees used. FastMulRFS (Molloy and Warnow, 2020) sometimes had high error rates when rates of ILS were high, a result that has been found in previous studies on the accuracy of this method (Zhang et al., 2020). FastMulRFS is also the only method employed here that has not been proven to be statistically consistent under the multispecies coalescent model, in which ILS is the driving forces behind incongruence.

In the cases where both ILS and GDL acted, the performance of the five methods was hardly affected by the type of dataset used (ALL, ONE, ONLY, ORTHOLOGS). Within the range of simulation parameters and datasets analyzed here, our results imply that running these methods on data with paralogs will produce species tree topologies at least as accurate as those using single-copy orthologs alone. This is especially important for datasets with a large number of species or high GDL rates.

When the methods were run on the locus tree data, where ILS does not play a role and the data consist of many gene families with multiple copies, the methods produced very accurate species trees. When as few as ten gene trees were used, error rates were elevated in datasets including paralogs (fig. S9). However, with more than ten genes, GDL alone did not appear to affect species tree inference under our simulation conditions. This further demonstrates that GDL has very little effect on the performance of these methods.

While at first it may be surprising that these methods performed very well in terms of accuracy, the majority of signal in any input gene tree reflects species relationships. Gene duplication—if random across the species tree—simply adds noise to the data, while at the same time often doubling the amount of information on the relationships among species carrying an extra gene copy. Similarly, gene loss does not positively mislead these methods, leading to accurate reconstructions of the species tree. Nevertheless, upon close inspection, some of these results are not intuitive, especially for the maximum pseudo-likelihood inference. `InferNetwork_MPL` makes direct use of the MSC, whose assumptions are clearly violated in all data sets except when the GDL rates are set to 0, whereas all other methods are summary methods that make no direct use of the MSC. Consequently, one would have expected that `InferNetwork_MPL` would be very sensitive to the presence of paralogs in the data, while the others were less so. However, we largely did not observe this behavior (but see discussion of the fungal tree below). Using methods designed specifically to deal with duplication and loss (ASTRAL-Pro and FastMulRFS) also did not lead to lower error rates. In the case of ASTRAL-Pro, we find performance similar to ASTRAL, as expected since both are statistically consistent under GDL and ILS (Legried et al., 2020; Zhang et al., 2020).

In practice, gene trees are estimated from sequence data and can be erroneous. Error in the gene tree estimates, rather than ILS, could explain much of the heterogeneity observed in phylogenomic analyses, especially at deeper nodes in a species tree (Scornavacca and Galtier, 2017). We showed the gene tree estimation error can indeed impact species tree inference significantly, and that the level of its impact is similar to that of ILS, if not larger. The results from simulations including gene tree error (and from the biological datasets) should be considered the most realistic. However, as more gene trees are used, regardless of levels of ILS or GDL, species tree accuracy increased

In analyses of two biological datasets where a species tree has been inferred using hundreds or thousands of loci, we found high accuracy of the methods using paralogs. All methods accurately inferred the published fly species tree. For the fungal species tree, no methods inferred the species tree we initially assumed to be true, which is originally based on a concatenated analysis of 706 single-copy genes (Butler et al., 2009). All methods, applied to all datasets, disagreed with this published tree with respect to the relative positions of *C. glabrata* and *S. castellii* (Fig. 7). Interestingly, the position of *S. castellii* in Butler et al. (2009) was constrained prior to tree search based on several rare genomic changes; an unconstrained search produced a topology consistent with the one found here. Shen et al. (2016), using a dataset of 1,233 single-copy orthologs, could not confidently determine the relationships among these species. Here, by more than doubling the number of gene trees, we find a species tree with a local posterior probability of 1.0 for the topology shown in Fig 7. Furthermore, the results of Shen et al. (2016) support the placement of *K. lactis* found here. The only sets of relationships that appears to differ with up-to-date fungal phylogenies are the ones inferred by `InferNetwork_MPL`(ALL) and `InferNetwork_MPL`(ONLY). This may be because `InferNetwork_MPL` explicitly models data according to the MSC.

All of our results point to a clear message: Under the conditions of our simulations and on the two biological datasets used here, running species tree inference methods intended to deal with ILS on gene trees with paralogs yields highly accurate results. This conclusion is powerful for at least two reasons. First, it implies that orthology assignment and paralogy removal are not necessary for running gene tree-based species tree inference; simply treating all copies as different individuals or randomly selecting a single copy would yield very accurate species tree topologies. Second, in many practical cases, too few single-copy genes are available to ensure good performance of species tree inference from those data alone. In these cases, our results suggest a ready source of more phylogenetic signal. Summary methods that do not explicitly use the MSC model (i.e., ASTRAL, ASTRAL-Pro, FastMulRFS, and NJ_st_) are expected to be more robust in the presence of GDL than methods that explicitly use the model—some of these methods have even been found to be statistically consistent under a model of GDL and ILS (Legried et al., 2020; Markin and Eulenstein, 2020).

While our study focused on the accuracy of the inferred species tree topology, using paralogs for inference would clearly have an impact on the estimated branch lengths of the species tree. In particular, under the ALL setting, there could be much more incongruence due to the large number of lineages, and, consequently, methods that use incongruence to estimate branch lengths would give values that are shorter than they truly are. For this reason, branch lengths inferred by such methods should not be used. Instead, an alternative approach is needed. The results of our analyses point to the following potential approach for inferring accurate species trees (topologies and branch lengths) by utilizing as much of the phylogenomic data as possible:

1. Use all available gene trees as input, whether or not they are single-copy in all species.
2. Feed all gene trees to a gene tree-based method to obtain a species tree topology.
3. Using a smaller subset of truly single-copy genes, and fixing the species tree topology obtained from Step (2), optimize the branch lengths of the species tree.

For Steps (1) and (2), one option is to repeat the random sampling of single copies from each species used to generate multiple “ONE” datasets. Then, these inferred species trees could be scored under some criterion that combines the MSC with a model of gene duplication/loss. This would overcome the issue of fixing a single species tree as input to Step (3), and avoids searching species tree space while evaluating a likelihood function that is very complex and computationally very demanding to compute. As an alternative to using only single-copy orthologs in Step (3), one could also use a statistical model that combines the MSC and GDL models (e.g., Rasmussen and Kellis, 2012). Such methods allow for paralogy detection and orthology assignment, conditional on the fixed species tree (or species trees), by using a more detailed evolutionary model and the full signal in the sequence data. For example, the orthology assignment could be “integrated out” or sampled, depending on the desired outcomes of the analysis. Unfortunately, while full Bayesian methods exist that model GDL alone (Boussau et al., 2013) or that model ILS alone (Ogilvie et al., 2017), none that we know of can model both.

## CONCLUSIONS

In this paper we set out to study how gene tree-based species tree inference would perform on data with paralogs. The motivation for exploring this question was two-fold. First, as methods for dealing with incongruence due to ILS have become commonplace, and as practitioners are almost never certain that their data contain no paralogs, it is important to understand the effect of hidden paralogy on the quality of the inference. Second, as larger phylogenomic datasets become available, insistence on single-copy genes would mean throwing away most of the data and potentially keeping a number of loci that may be inadequate for suitably complex species tree inference methods to perform well. We investigated this question through a combination of simulations and biological data analyses. Our results show that gene tree-based inference is robust to the presence of paralogs in the data, at least under the simulation conditions and on the empirical datasets we investigated.

Our results highlight the issue that gene tree-based inference could result in very accurate species trees even when ILS is not a factor or not the only factor. This finding implies that orthology detection and restricting data to single-copy genes as a requirement for employing gene tree-based inference can be mostly eliminated, thus making use of as much of the data as possible (cf. Smith and Hahn, 2021b). In particular, for very large datasets (in terms of the number of species), eliminating all but single-copy genes might leave too few loci for the species tree to be inferred accurately. Our findings show that this data exclusion could be an unnecessary practice. It is important to note however, that our results do not apply to concatenated analyses, and in such cases the presence of paralogs may indeed have a large, negative effect (Brown and Thomson, 2016). Species tree inference from a concatenation of the sequences with gene families is challenging in the presence of paralogs for at least two reasons. First, when gene families have different numbers of copies across species, the concatenated alignment will have very large gaps. Second, correct orthology detection is still required, so that orthologous gene copies are placed in correct correspondence across the multiple genomes in the concatenated alignment. This issue is very important to examine so as to avoid aligning non-orthologous sequences in the concatenated data set.

In our simulations, we generated gene families under a neutral model and with GDL rates that were the same across all families. It is well known that the functional implications of gene duplication and the ways in which they are fixed and maintained in the genome result in much more complex scenarios than those captured in our simulations (Hahn, 2009; Innan and Kondrashov, 2010). However, analyses of the two biological datasets yield results with very similar trends to those observed in our simulations.

Finally, while we did not discuss or incorporate gene flow in our study, it is possible that all three processes—ILS, GDL, and gene flow—are simultaneously involved in the evolution of some clades. Studies of the robustness of gene tree-based species tree inference under some models of gene flow exist (Roch and Snir, 2012; Steel et al., 2013; Davidson et al., 2015; SolÍs-Lemus et al., 2016; Zhu et al., 2016; Long and Kubatko, 2018), but, to the best of our knowledge, such studies under scenarios that incorporate all the aforementioned processes do not exist yet. It is important to highlight, as well, that great strides have been made in developing methods for phylogenetic network inference in the presence of ILS (Elworth et al., 2019), but no probabilistic methods currently incorporate gene duplication and loss (see Li et al. (2020) for a very interesting alternative approach). We believe methods along the lines described in the previous section could be promising for accurate and scalable phylogenomic inferences without sacrificing much of the data.

## Supporting information

SIupplementary Material

## SUPPLEMENTARY MATERIAL

Supplementary material, including data files and online-only appendices, can be found in the Dryad data repository at

## References

Arvestad, L., J. Lagergren, and B. Sennblad. 2009. The gene evolution model and computing its associated probabilities. Journal of the ACM 56:7.

Boussau, B., G. J. Szöllősi, L. Duret, M. Gouy, E. Tannier, and V. Daubin. 2013. Genome-scale coestimation of species and gene trees. Genome Research 23:323–330.

Brown, J. M. and R. C. Thomson. 2016. Bayes factors unmask highly variable information content, bias, and extreme influence in phylogenomic analyses. Systematic Biology 66:517–530.

Bryant, D. and M. W. Hahn. 2020. The concatenation question. Pages 3.4:1–3.4:23 in Phylogenetics in the Genomic Era (C. Scornavacca, F. Delsuc, and N. Galtier, eds.). No commercial publisher — Authors open access book.

Butler, G., M. D. Rasmussen, M. F. Lin, M. A. Santos, S. Sakthikumar, C. A. Munro, E. Rheinbay, M. Grabherr, A. Forche, J. L. Reedy, et al. 2009. Evolution of pathogenicity and sexual reproduction in eight *candida* genomes. Nature 459:657–662.

Davidson, R., P. Vachaspati, S. Mirarab, and T. Warnow. 2015. Phylogenomic species tree estimation in the presence of incomplete lineage sorting and horizontal gene transfer. BMC Genomics 16:S1.

Degnan, J. H. and N. A. Rosenberg. 2009. Gene tree discordance, phylogenetic inference and the multispecies coalescent. Trends in Ecology & Evolution 24:332–340.

Doolittle, W. F. and J. R. Brown. 1994. Tempo, mode, the progenote, and the universal root. Proceedings of the National Academy of Sciences 91:6721–6728.

Du, P. and L. Nakhleh. 2018. Species tree and reconciliation estimation under a duplication-loss-coalescence model. Proceedings of the 9th ACM Conference on Bioinformatics, Computational Biology, and Health Informatics Pages 376–385.

Elworth, R. L., H. A. Ogilvie, J. Zhu, and L. Nakhleh. 2019. Advances in computational methods for phylogenetic networks in the presence of hybridization. Pages 317–360 in Bioinformatics and Phylogenetics (T. Warnow, ed.). Springer.

Emms, D. and S. Kelly. 2018. STAG: Species tree inference from all genes. bioRxiv Page 267914.

Guindon, S. and O. Gascuel. 2003. A simple, fast, and accurate algorithm to estimate large phylogenies by maximum likelihood. Systematic Biology 52:696–704.

Hahn, M. W. 2009. Distinguishing among evolutionary models for the maintenance of gene duplicates. Journal of Heredity 100:605–617.

Hahn, M. W., M. V. Han, and S.-G. Han. 2007. Gene family evolution across 12 drosophila genomes. PLOS Genetics 3:e197.

Hudson, R. R. 1983. Testing the constant-rate neutral allele model with protein sequence data. Evolution 37:203–217.

Innan, H. and F. Kondrashov. 2010. The evolution of gene duplications: classifying and distinguishing between models. Nature Reviews Genetics 11:97–108.

Knowles, L. L. and L. S. Kubatko. 2011. Estimating species trees: practical and theoretical aspects. John Wiley and Sons.

Koonin, E. V. 2005. Orthologs, paralogs, and evolutionary genomics. Annu. Rev. Genet. 39:309–338.

Lang, G. I. and A. W. Murray. 2008. Estimating the per-base-pair mutation rate in the yeast *saccharomyces cerevisiae*. Genetics 178:67–82.

Legried, B., E. K. Molloy, T. Warnow, and S. Roch. 2020. Polynomial-time statistical estimation of species trees under gene duplication and loss. Journal of Computational Biology ahead of print:cmb.2020.0424.

Li, L., C. J. Stoeckert Jr., and D. S. Roos. 2003. OrthoMCL: Identification of Ortholog groups for eukaryotic genomes. Genome Research 13:2178–2189.

Li, Q., C. Scornavacca, N. Galtier, and Y.-B. Chan. 2020. The multilocus multispecies coalescent: a flexible new model of gene family evolution. Systematic Biology Syaa084.

Liu, L., Z. Xi, S. Wu, C. C. Davis, and S. V. Edwards. 2015. Estimating phylogenetic trees from genome-scale data. Annals of the New York Academy of Sciences 1360:36–53.

Liu, L. and L. Yu. 2011. Estimating species trees from unrooted gene trees. Systematic Biology 60:661–667.

Liu, L., L. Yu, and S. V. Edwards. 2010. A maximum pseudo-likelihood approach for estimating species trees under the coalescent model. BMC Evolutionary Biology 10:302.

Liu, L., L. L. Yu, L. Kubatko, D. K. Pearl, and S. V. Edwards. 2009. Coalescent methods for estimating phylogenetic trees. Molecular Phylogenetics and Evolution 53:320–328.

Long, C. and L. Kubatko. 2018. The effect of gene flow on coalescent-based species-tree inference. Systematic biology 67:770–785.

Maddison, W. P. 1997. Gene trees in species trees. Systematic Biology 46:523–536.

Mallo, D., L. de Oliveira Martins, and D. Posada. 2015. SimPhy: phylogenomic simulation of gene, locus, and species trees. Systematic Biology 65:334–344.

Markin, A. and O. Eulenstein. 2020. Quartet-based inference methods are statistically consistent under the unified duplication-loss-coalescence model. arXiv preprint arXiv:2004.04299.

Mirarab, S., R. Reaz, M. S. Bayzid, T. Zimmermann, M. S. Swenson, and T. Warnow. 2014. ASTRAL: genome-scale coalescent-based species tree estimation. Bioinformatics 30:i541–i548.

Molloy, E. K. and T. Warnow. 2020. FastMulRFS: fast and accurate species tree estimation under generic gene duplication and loss models. Bioinformatics 36:i57–i65.

Nakhleh, L. 2013. Computational approaches to species phylogeny inference and gene tree reconciliation. Trends in Ecology and Evolution 28:719–728.

Nguyen, L.-T., H. A. Schmidt, A. von Haeseler, and B. Q. Minh. 2014. IQ-TREE: a fast and effective stochastic algorithm for estimating maximum-likelihood phylogenies. Molecular Biology and Evolution 32:268–274.

Ogilvie, H. A., R. R. Bouckaert, and A. J. Drummond. 2017. StarBEAST2 brings faster species tree inference and accurate estimates of substitution rates. Molecular Biology and Evolution 34:2101–2114.

Pollard, D. A., V. N. Iyer, A. M. Moses, and M. B. Eisen. 2006. Widespread discordance of gene trees with species tree in *drosophila*: evidence for incomplete lineage sorting. PLOS Genetics 2:e173.

Rambaut, A. and N. C. Grassly. 1997. Seq-Gen: an application for the Monte Carlo simulation of DNA sequence evolution along phylogenetic trees. Bioinformatics 13:235–238.

Rannala, B. and Z. Yang. 2003. Bayes estimation of species divergence times and ancestral population sizes using dna sequences from multiple loci. Genetics 164:1645–1656.

Rasmussen, M. D. and M. Kellis. 2011. A Bayesian approach for fast and accurate gene tree reconstruction. Molecular Biology and Evolution 28:273–290.

Rasmussen, M. D. and M. Kellis. 2012. Unified modeling of gene duplication, loss, and coalescence using a locus tree. Genome Research 22:755–765.

Robinson, D. F. and L. R. Foulds. 1981. Comparison of phylogenetic trees. Mathematical Biosciences 53:131–147.

Roch, S. and S. Snir. 2012. Recovering the tree-like trend of evolution despite extensive lateral genetic transfer: a probabilistic analysis. Journal of Computational Biology 20:93–112.

Sawyer, S. A. and D. L. Hartl. 1992. Population genetics of polymorphism and divergence. Genetics 132:1161–1176.

Schrider, D. R., D. Houle, M. Lynch, and M. W. Hahn. 2013. Rates and genomic consequences of spontaneous mutational events in Drosophila melanogaster. Genetics 194:937–954.

Scornavacca, C. and N. Galtier. 2017. Incomplete lineage sorting in mammalian phylogenomics. Systematic Biology 66:112–120.

Shen, X.-X., X. Zhou, J. Kominek, C. P. Kurtzman, C. T. Hittinger, and A. Rokas. 2016. Reconstructing the backbone of the Saccharomycotina yeast phylogeny using genome-scale data. G3: Genes, Genomes, Genetics 6:3927–3939.

Smith, M. L. and M. W. Hahn. 2021a. The frequency and topology of pseudoorthologs. bioRxiv Page 10.1101/2021.02.17.431499.

Smith, M. L. and M. W. Hahn. 2021b. New approaches for inferring phylogenies in the presence of paralogs. Trends in Genetics 37:174–187.

SolÍs-Lemus, C., M. Yang, and C. Ané. 2016. Inconsistency of species tree methods under gene flow. Systematic Biology 65:843–851.

Steel, M., S. Linz, D. Huson, and M. Sanderson. 2013. Identifying a species tree subject to random lateral gene transfer. Journal of Theoretical Biology 322:81–93.

Takahata, N. 1989. Gene genealogy in three related populations: consistency probability between gene and population trees. Genetics 122:957–966.

Than, C. and L. Nakhleh. 2009. Species tree inference by minimizing deep coalescences. PLoS Computational Biology 5:e1000501.

Thomas, G. W., E. Dohmen, D. S. Hughes, S. C. Murali, M. Poelchau, K. Glastad, C. A. Anstead, N. A. Ayoub, P. Batterham, M. Bellair, et al. 2020. Gene content evolution in the arthropods. Genome Biology 21:15.

Wen, D., Y. Yun, J. Zhu, and L. Nakhleh. 2018. Inferring phylogenetic networks using PhyloNet. Systematic Biology 67:735–740.

Yang, Y. and S. A. Smith. 2014. Orthology inference in nonmodel organisms using transcriptomes and low-coverage genomes: Improving accuracy and matrix occupancy for phylogenomics. Molecular Biology and Evolution 31:3081–3082.

Yu, Y., J. Dong, K. J. Liu, and L. Nakhleh. 2014. Maximum likelihood inference of reticulate evolutionary histories. Proceedings of the National Academy of Sciences 111:16448–16453.

Yu, Y. and L. Nakhleh. 2015. A maximum pseudo-likelihood approach for phylogenetic networks. BMC Genomics 16:S10.

Yu, Y., T. Warnow, and L. Nakhleh. 2011. Algorithms for MDC-based multi-locus phylogeny inference: beyond rooted binary gene trees on single alleles. Journal of Computational Biology 18:1543–1559.

Zhang, B. and Y.-C. Wu. 2017. Coestimation of gene trees and reconciliations under a duplication-loss-coalescence model. Pages 196–210 in International Symposium on Bioinformatics Research and Applications Springer.

Zhang, C., M. Rabiee, E. Sayyari, and S. Mirarab. 2018. ASTRAL-III: polynomial time species tree reconstruction from partially resolved gene trees. BMC Bioinformatics 19:153.

Zhang, C., C. Scornavacca, E. K. Molloy, and S. Mirarab. 2020. ASTRAL-Pro: quartet-based species-tree inference despite paralogy. Molecular Biology and Evolution 37:3292–3307.

Zhu, J., Y. Yu, and L. Nakhleh. 2016. In the light of deep coalescence: revisiting trees within networks. BMC Bioinformatics 17:415.

