## Supplementary material for "Species Tree Inference on Data with Paralogs is Accurate Using Methods Intended to Deal with Incomplete Lineage Sorting": SIupplementary Material

- Characteristics of the 16-taxon simulated yeast data sets of population size $10^7$ are shown in Fig. S1 (a), (c) and (e).
- Characteristics of the 12-taxon simulated fly data sets of population size $10^6$ are shown in Fig. S1 (b), (d) and (f).
- Accuracy of inferred species trees on the 16-taxon simulated yeast data sets of population size $10^7$ from true gene trees, true locus trees, estimated gene trees, and estimated locus trees, are shown in Fig. S4—S7.
- Gene tree and locus tree estimation error on the 16-taxon simulated yeast data sets are shown in Fig. S2.
- Gene tree and locus tree estimation error on the 12-taxon simulated fly data sets are shown in Fig. S3.
- Accuracy of inferred species trees on the 16-taxon simulated yeast data sets of population size $10^7$ from true gene trees, true locus trees, estimated gene trees, and estimated locus trees, are shown in Fig. S4—S7.
- Accuracy of inferred species trees on the 12-taxon simulated fly data sets from true gene trees, true locus trees, estimated gene trees, and estimated locus trees, are shown in Fig. S8—S11.

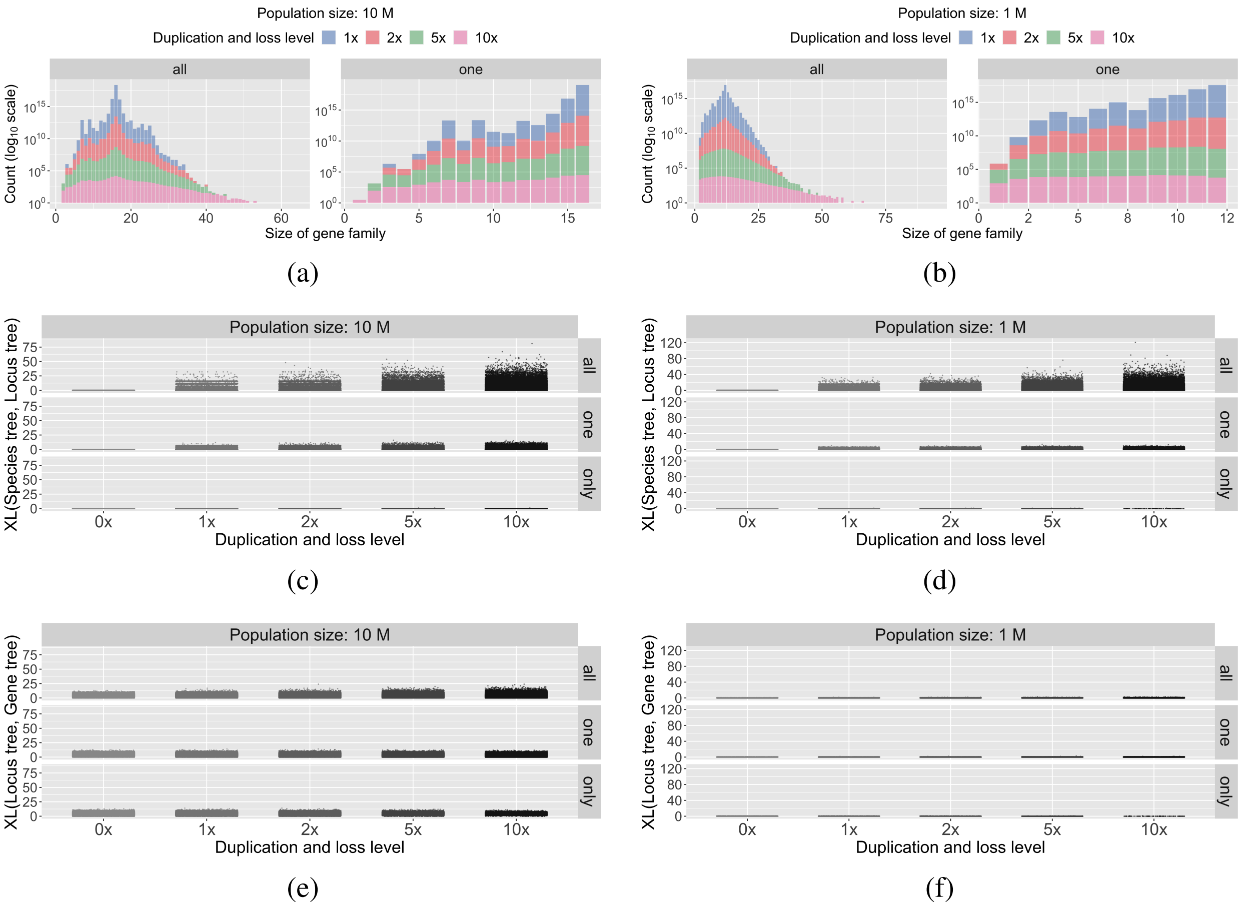


Figure S1: Characteristics of the simulated data under different settings of the duplication/loss rates and tree topologies. The duplication/loss rates are denoted by the rate multiplier (0x, 1x, 2x, 5x and 10x), where 1x is the rate found in nature for the clade represented by each species tree topology (see Methods). (a-b) Distribution of the total number of gene copies in individual gene families in the 16-taxon and 12-taxon data sets, respectively. (c-d) Scatter plots of XL(Species tree, Locus tree), the number of extra lineages when reconciling the true locus trees with the true species tree, for the 16-taxon and 12-taxon data sets, respectively. These plots therefore represent the effects of GDL alone. (e-f) Scatter plots of XL(Locus tree, Gene tree), the number of extra lineages when reconciling the true gene trees with the true locus tree, for the 16-taxon and 12-taxon data sets, respectively. These plots therefore represent the effects of ILS alone, though note that higher rates of GDL allow there to be more gene tree branches on which ILS can act.

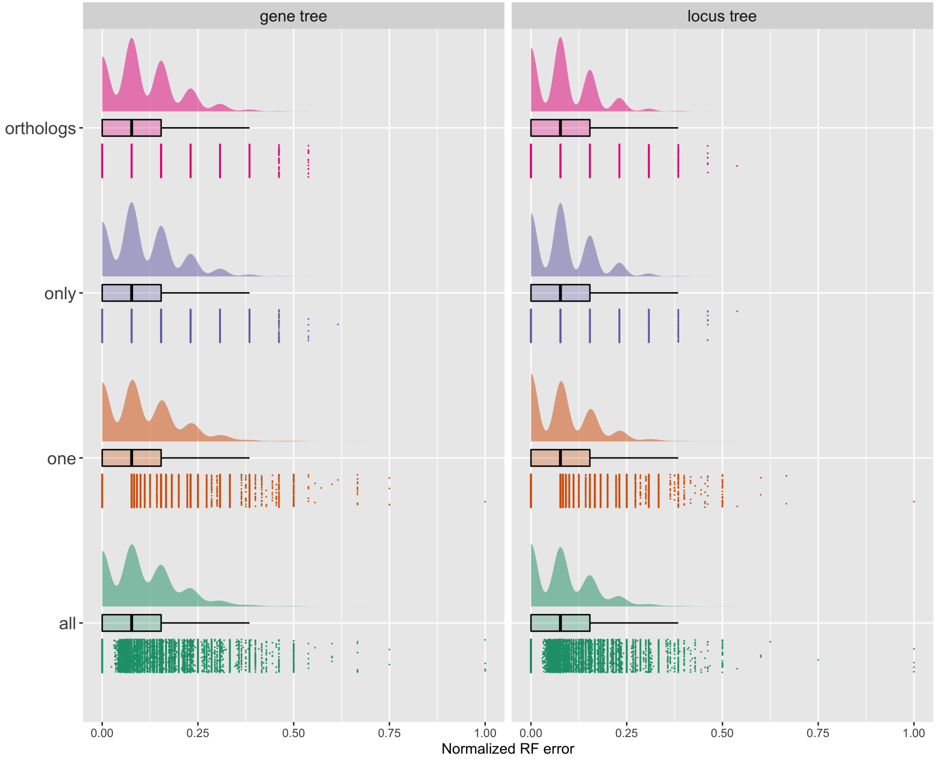


Figure S2: The normalized RF distances between the true and estimated gene trees as well as the true and estimated locus trees for the 16-taxon simulated yeast data sets. Gene trees and locus trees were inferred from sequence data using IQ-TREE.

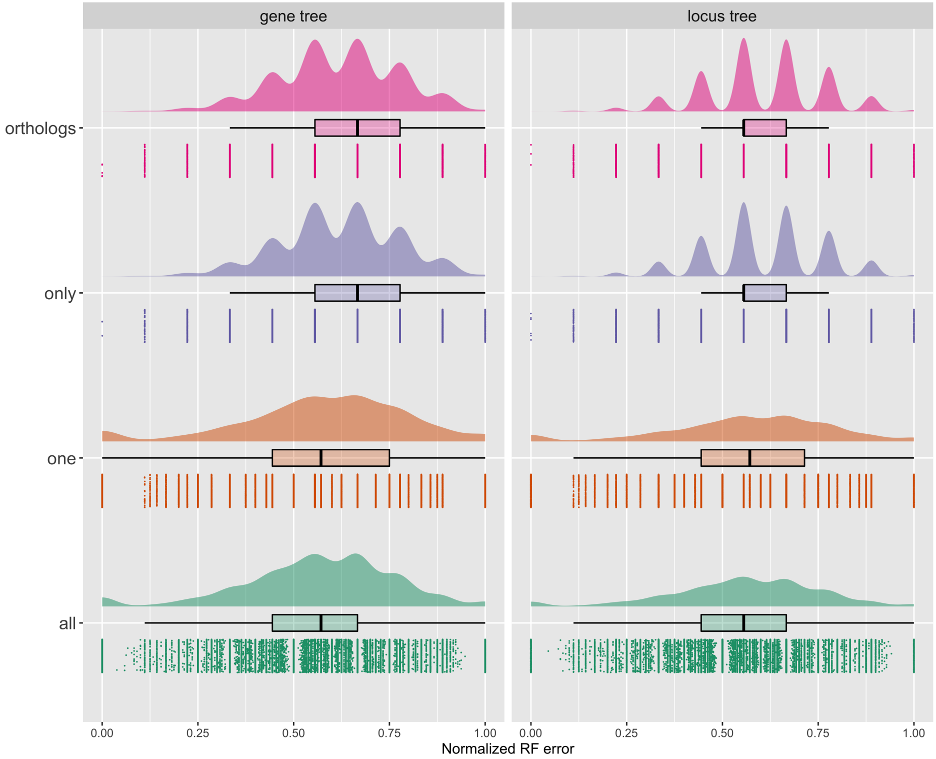


Figure S3: The normalized RF distances between the true and estimated gene trees as well as the true and estimated locus trees for the 12-taxon simulated fly data sets. Gene trees and locus trees were inferred from sequence data using IQ-TREE.

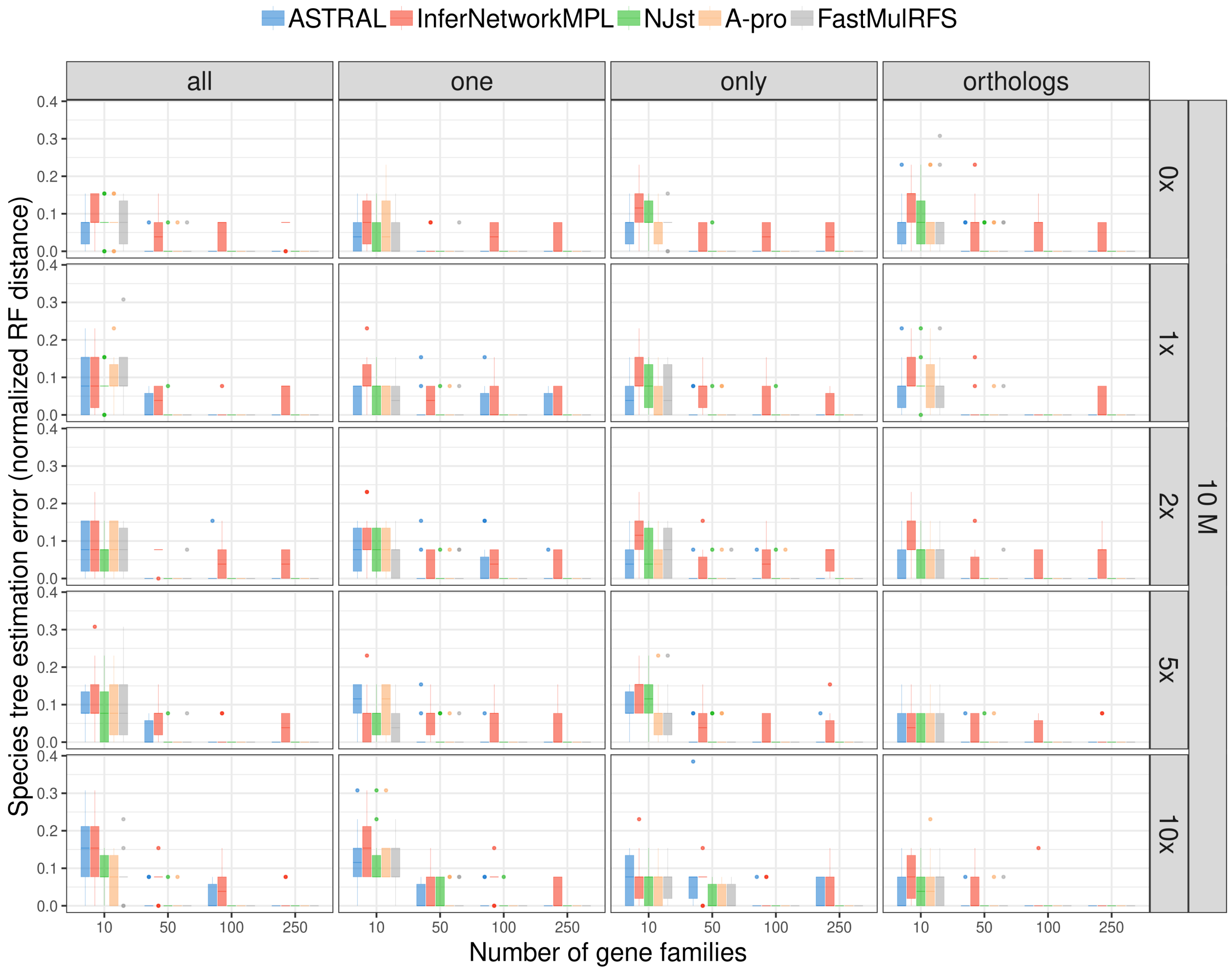


Figure S4: The normalized RF distances between the true species tree and the ones inferred from true gene trees. These simulations therefore include the effects of both ILS and GDL (but no gene tree estimation error). The five inference methods used are ASTRAL, `InferNetwork_MPL`, $NJ_{st}$, ASTRAL-Pro ("A-pro"), and FastMulRFS), and data is simulated on the 16-taxon tree with a population size of $10^7$ and varying GDL rates. The duplication/loss rates are denoted by the rate multiplier (0x, 1x, 2x, 5x and 10x), where 1x is the rate estimated in nature for fungi. Each row corresponds to a combination of population size and GDL rates. The X-axis in each panel represents the number of gene families used and the Y-axis represents the normalized RF distance.

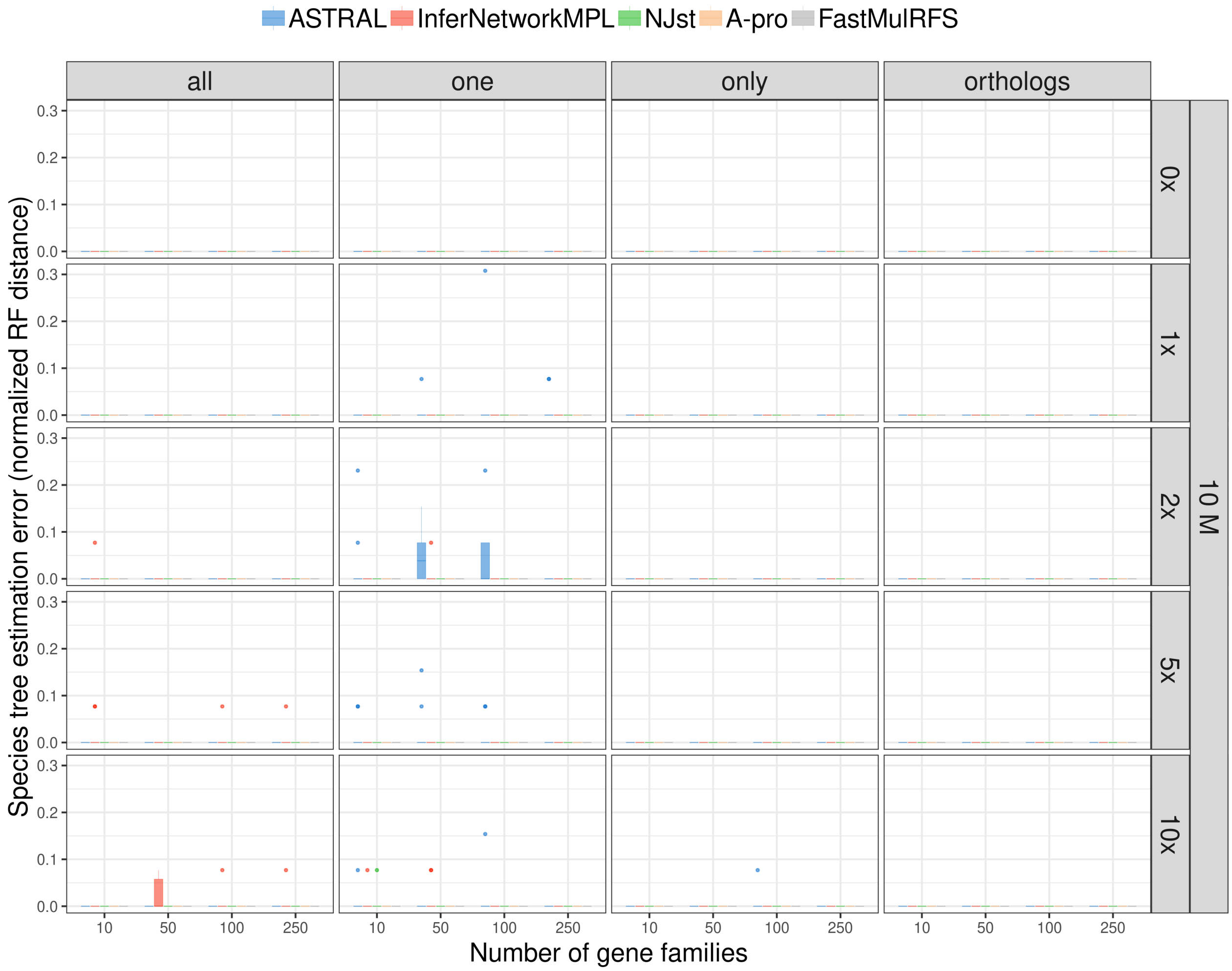


Figure S5: The normalized RF distances between the true species tree and the ones inferred from true locus trees. These simulations therefore include the effects of GDL only (no ILS or gene tree estimation error). The five inference methods used are ASTRAL, `InferNetwork_MPL`, $NJ_{st}$, ASTRAL-Pro ("A-pro"), and FastMulRFS), and data is simulated on the 16-taxon tree with a population size of $10^7$ and varying GDL rates. The duplication/loss rates are denoted by the rate multiplier (0x, 1x, 2x, 5x and 10x), where 1x is the rate estimated in nature for fungi. Each row corresponds to a combination of population size and GDL rates. The X-axis in each panel represents the number of gene families used and the Y-axis represents the normalized RF distance.

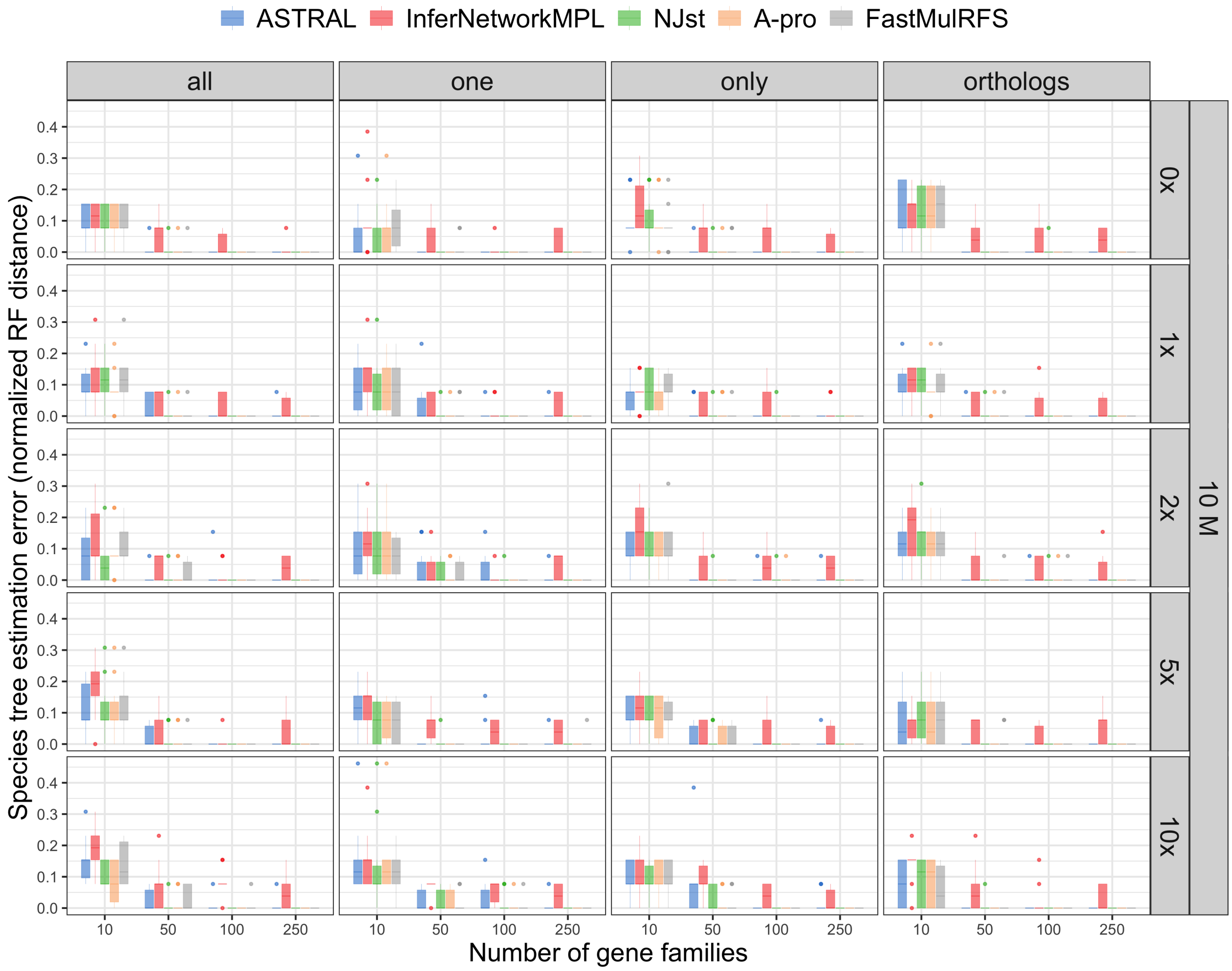


Figure S6: The normalized RF distances between the true species tree and the ones inferred from estimated gene trees. These simulations therefore include the effects of GDL, ILS, and gene tree estimation error. The five inference methods used are ASTRAL, `InferNetwork_MPL`, $NJ_{st}$, ASTRAL-Pro ("A-pro"), and FastMulRFS), and data is simulated on the 16-taxon tree with a population size of $10^7$ and varying GDL rates. The duplication/loss rates are denoted by the rate multiplier (0x, 1x, 2x, 5x and 10x), where 1x is the rate estimated in nature for fungi. Each row corresponds to a combination of population size and GDL rates. The X-axis in each panel represents the number of gene families used and the Y-axis represents the normalized RF distance.

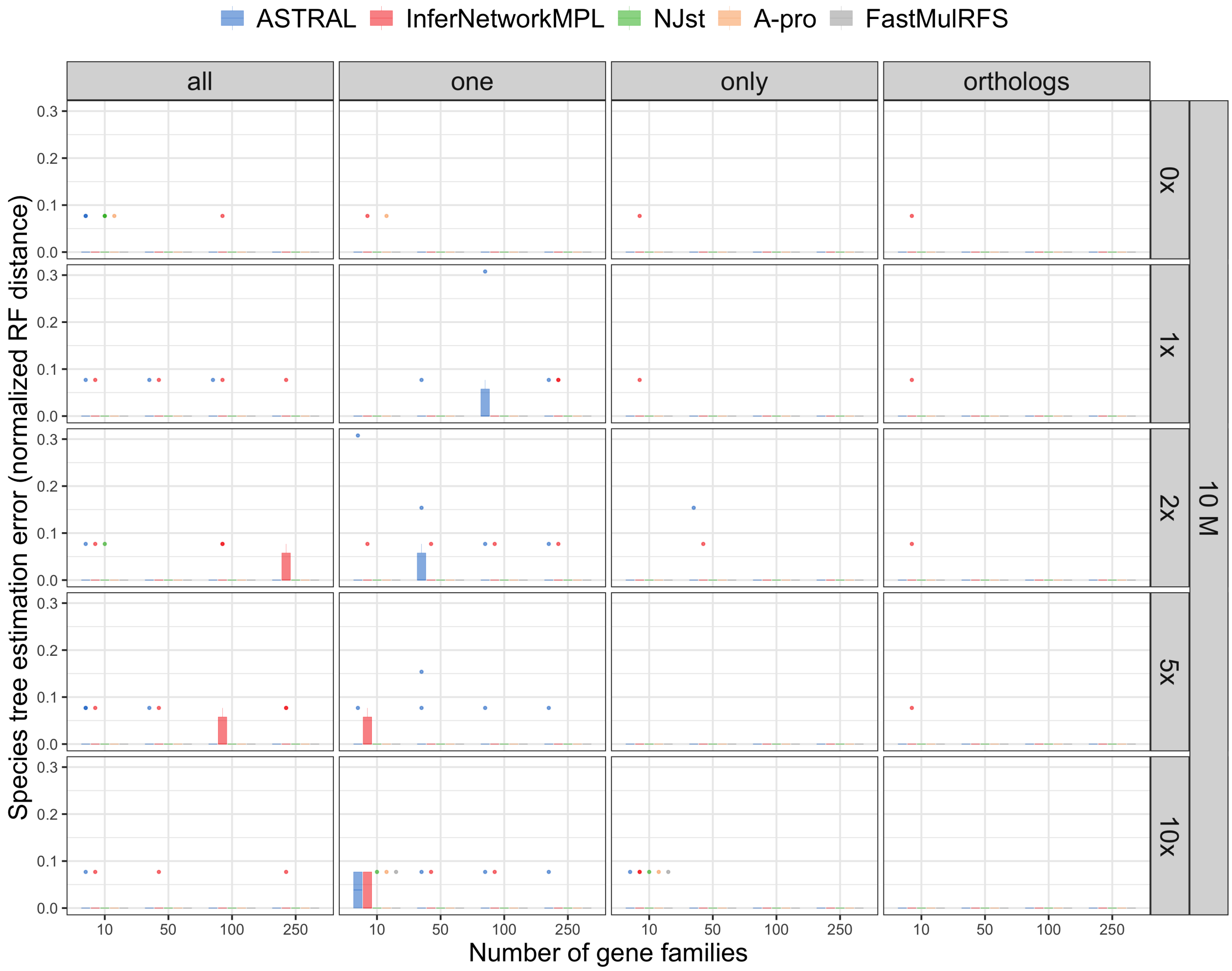


Figure S7: The normalized RF distances between the true species tree and the ones inferred from the estimated locus trees. These simulations therefore include the effects of GDL and gene tree estimation error (no ILS). The five inference methods used are ASTRAL, `InferNetwork_MPL`, $NJ_{st}$, ASTRAL-Pro ("A-pro"), and FastMulRFS), and data is simulated on the 16-taxon tree with a population size of $10^7$ and varying GDL rates. The duplication/loss rates are denoted by the rate multiplier (0x, 1x, 2x, 5x and 10x), where 1x is the rate estimated in nature for fungi. Each row corresponds to a combination of population size and GDL rates. The X-axis in each panel represents the number of gene families used and the Y-axis represents the normalized RF distance.

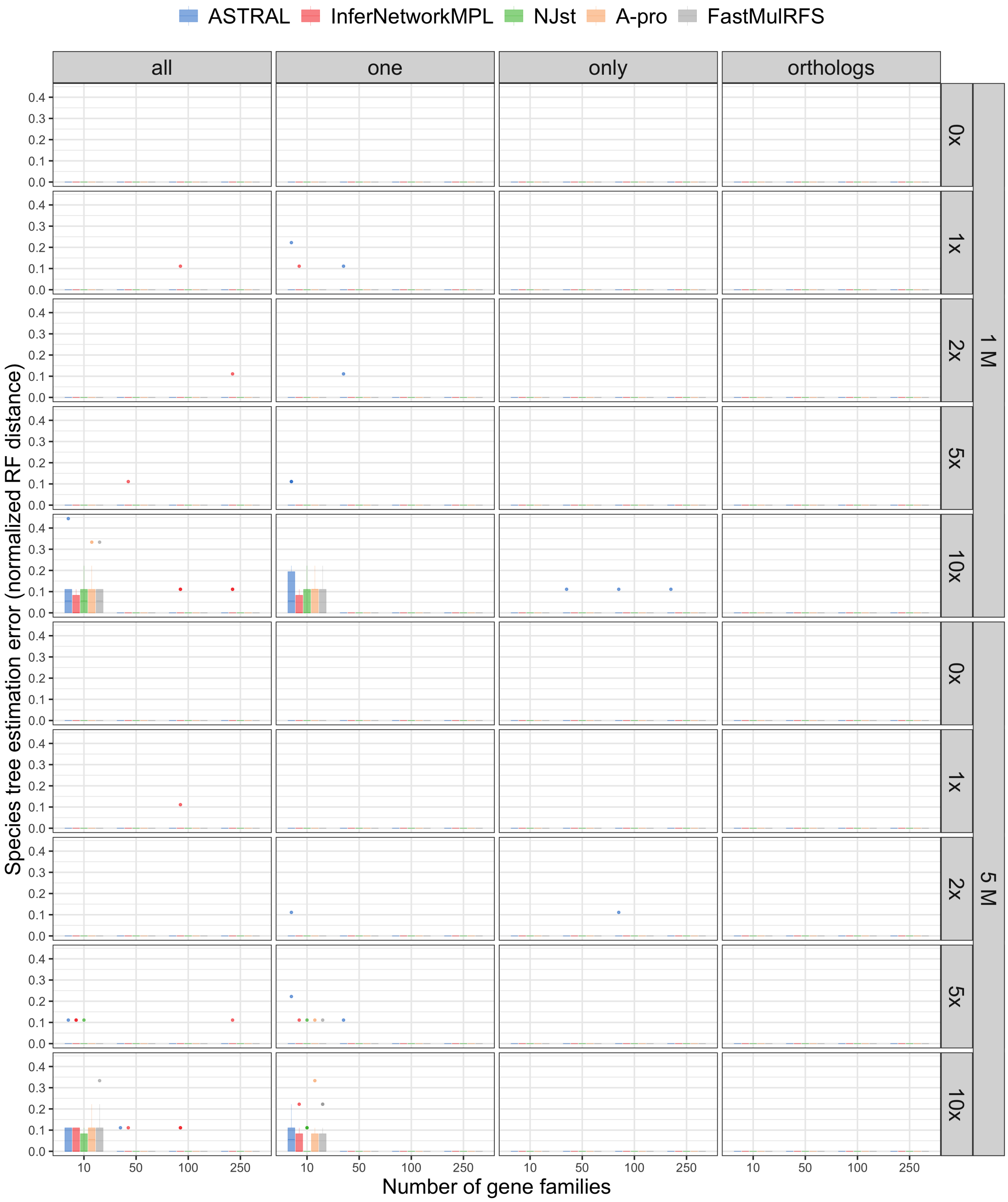


Figure S8: The normalized RF distances between the true species tree and the ones inferred from true gene trees. These simulations therefore include the effects of both ILS and GDL (but no gene tree estimation error). The five inference methods used are ASTRAL, `InferNetwork_MPL`, $NJ_{st}$, ASTRAL-Pro ("A-pro"), and FastMulRFS), and data is simulated on the 12-taxon tree under different GDL rates and population sizes. The duplication/loss rates are denoted by the rate multiplier (0x, 1x, 2x, 5x and 10x), where 1x is the rate estimated in nature for fly. Each row corresponds to a combination of population size and GDL rates. The X-axis in each panel represents the number of gene families used and the Y-axis represents the normalized RF distance.

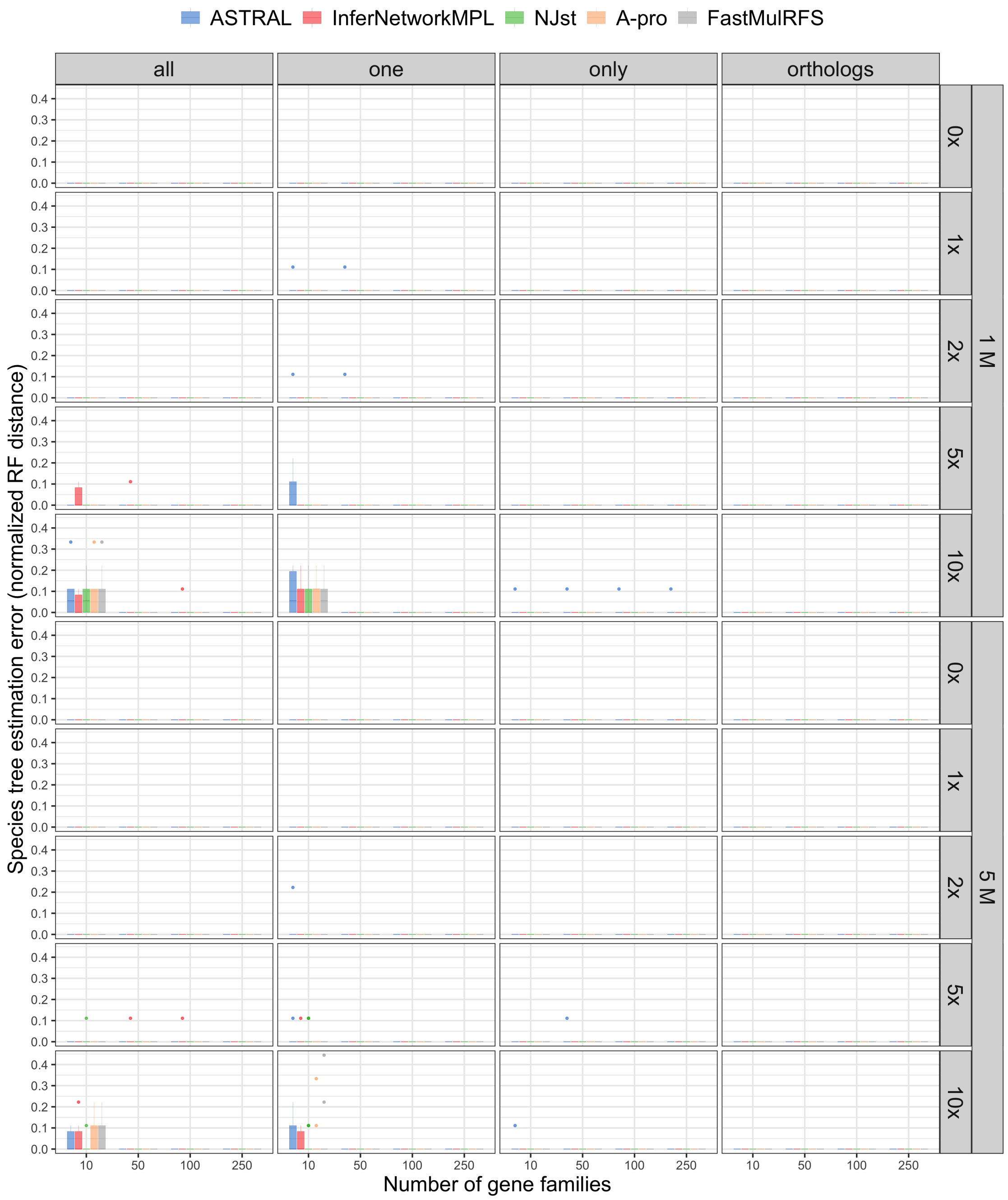


Figure S9: The normalized RF distances between the true species tree and the ones inferred from true locus trees. These simulations therefore include the effects of GDL only (no ILS or gene tree estimation error). The five inference methods used are ASTRAL, `InferNetwork_MPL`, $NJ_{st}$, ASTRAL-Pro ("A-pro"), and FastMulRFS), and data is simulated on the 12-taxon tree under different GDL rates and population sizes. The duplication/loss rates are denoted by the rate multiplier (0x, 1x, 2x, 5x and 10x), where 1x is the rate estimated in nature for fly. Each row corresponds to a combination of population size and GDL rates. The X-axis in each panel represents the number of gene families used and the Y-axis represents the normalized RF distance.

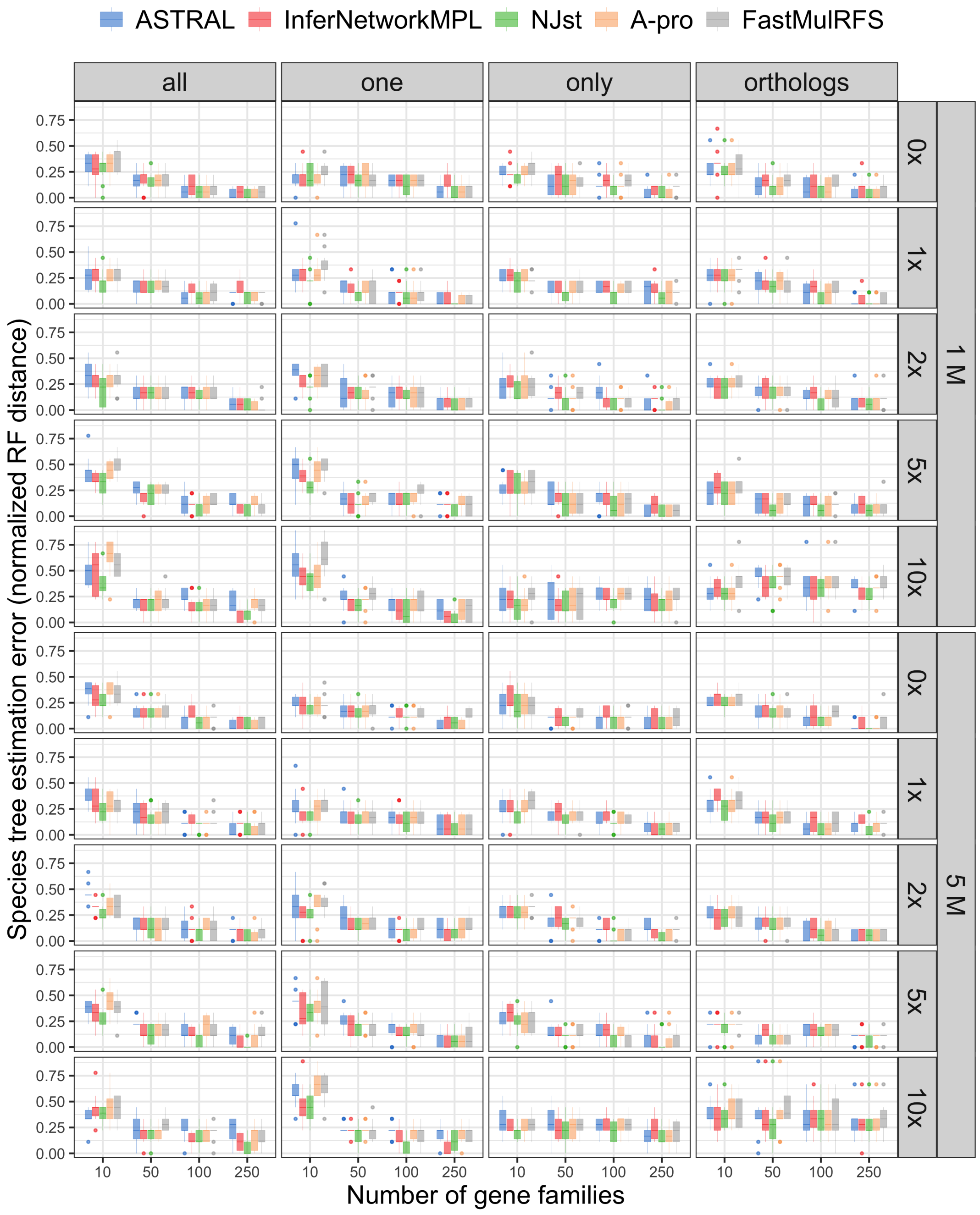


Figure S10: The normalized RF distances between the true species tree and the ones inferred from estimated gene trees. These simulations therefore include the effects of GDL, ILS, and gene tree estimation error. The five inference methods used are ASTRAL, `InferNetwork_MPL`, $NJ_{st}$, ASTRAL-Pro ("A-pro"), and FastMulRFS), and data is simulated on the 12-taxon tree under different GDL rates and population sizes. The duplication/loss rates are denoted by the rate multiplier (0x, 1x, 2x, 5x and 10x), where 1x is the rate estimated in nature for fly. Each row corresponds to a combination of population size and GDL rates. The X-axis in each panel represents the number of gene families used and the Y-axis represents the normalized RF distance.

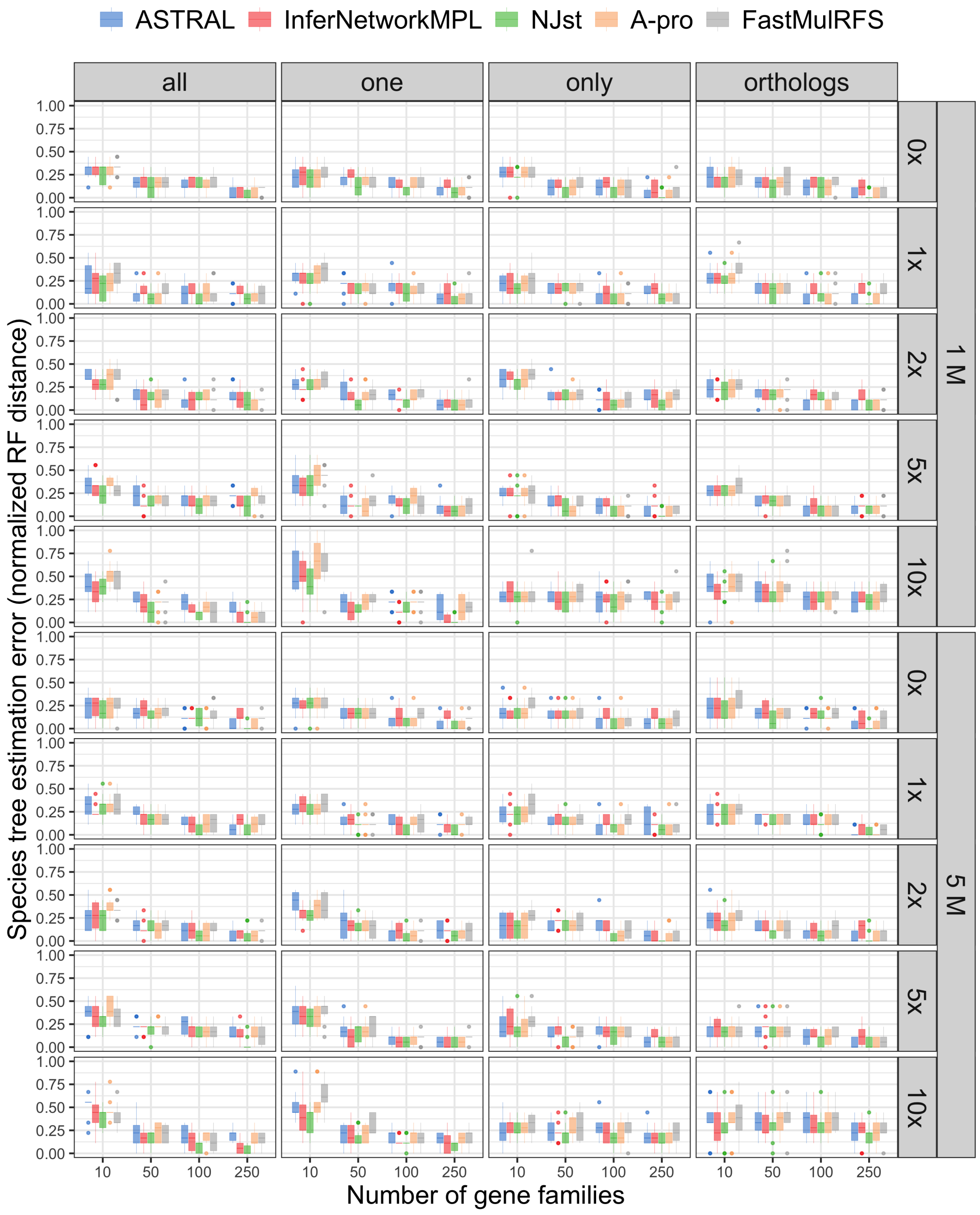


Figure S11: The normalized RF distances between the true species tree and the ones inferred from the estimated locus trees. These simulations therefore include the effects of GDL and gene tree estimation error (no ILS). The five inference methods used are ASTRAL, `InferNetwork_MPL`, $NJ_{st}$, ASTRAL-Pro ("A-pro"), and FastMulRFS), and data is simulated on the 12-taxon tree under different GDL rates and population sizes. The duplication/loss rates are denoted by the rate multiplier (0x, 1x, 2x, 5x and 10x), where 1x is the rate estimated in nature for fly. Each row corresponds to a combination of population size and GDL rates. The X-axis in each panel represents the number of gene families used and the Y-axis represents the normalized RF distance.
